# Graph analysis of looming-selective networks in the tectum, and its replication in a simple computational model

**DOI:** 10.1101/589887

**Authors:** Arseny S. Khakhalin

## Abstract

Looming stimuli evoke behavioral responses in most animals, yet the mechanisms of looming detection in vertebrates are poorly understood. Here we hypothesize that looming detection in the tectum may rely on spontaneous emergence of synfire chains: groups of neurons connected to each other in the same sequence in which they are activated during a loom. We then test some specific consequences of this hypothesis. First, we use high-speed calcium imaging to reconstruct functional connectivity of small networks within the tectum of Xenopus tadpoles. We report that reconstructed directed graphs are clustered and hierarchical, that their modularity increases in development, and that looming-selective cells tend to collect activation within these graphs. Second, we describe spontaneous emergence of looming selectivity in a computational developmental model of the tectum, governed by both synaptic and intrinsic plasticity, and driven by structured visual inputs. We show that synfire chains contribute to looming detection in the model; that structured inputs are critical for the emergence of selectivity, and that biological tectal networks follow most, but not all predictions of the model. Finally, we propose a conceptual scheme for understanding the emergence and fine-tuning of collision detection in developing aquatic animals.

## Introduction

Few sensory stimuli are as ill boding for the animal as a visual loom. A retinal projection that is small, but is quickly growing in size, may promise a painful collision, or a meeting with a predator, so it inherently calls for action: an avoidance maneuver, or a defensive response. Moreover, to be meaningful, looming detection has to be fast. Not surprisingly, it is described in virtually every animal that uses vision, from insects to primates (Pereira and Moita, 2016). And yet, while our understanding of looming detection in insects has recently improved (Rind et al., 2016; von Reyn et al., 2017), the mechanisms that underlie collision avoidance in vertebrates are still unclear.

For vertebrates, it is well established that looming detection primarily happens in a midbrain region known as superior colliculus in mammals, and optic tectum in all other clades (Frost and Sun, 2004; Liu et al., 2011; Khakhalin et al., 2014; Dunn et al., 2016). It is not known however how midbrain circuits perform the computations required for collision avoidance, and it is still unclear whether looming avoidance in vertebrates is innate (and so possibly hardwired), or whether it needs to be learned. Finally, to navigate, animals need to calculate not just *whether* a collision is about to happen, but *where* a looming stimulus is coming from. While it is known that the tectum harbors a retinotopic map of the visual field (McLaughlin et al., 2003; Ruthazer and Cline, 2004), there are several competing theories about how looming detectors may be distributed within this map (Frost and Sun, 2004). What calculations may in principle underlie looming detection? Across species, looming detection relies on a diverse set of mechanisms that include dimming detectors (Ishikane et al., 2005; Münch et al., 2009; Heap et al., 2018a), integration of opponent motion (Klapoetke et al., 2017), and competitive spike-frequency adaptation (Peron and Gabbiani, 2009; Fotowat et al., 2011). Even within a single clade of anuran amphibians (frogs), animals seem to employ at least two competing approaches to loom detection: a non-linear response to dimming-induced retinal oscillations (Baranauskas et al., 2012), and a rebound of recurrent activity during edge expansion (Khakhalin et al., 2014; Jang et al., 2016). Moreover, at least in some cases, competing mechanisms may lead to different motor responses, as described in insects (Card and Dickinson, 2008; Chan and Gabbiani, 2013), fish (Budick and O’Malley, 2000; Burgess and Granato, 2007; Portugues and Engert, 2009; Temizer et al., 2015; Bhattacharyya et al., 2017; Heap et al., 2018a), and tadpoles (Khakhalin et al., 2014).

It may seem puzzling that the brain would use several conflicting approaches to solve one practical problem, but this arrangement may make sense developmentally. Simple, crude ways of detecting dangerous stimuli can be used to train more sophisticated networks, to support efficient sensory analysis at later stages of development (Marblestone et al., 2016; Zador, 2019). We argue that young aquatic animals may use “hardwired” dimming receptors (Baranauskas et al., 2012; Heap et al., 2018a) both to avoid collisions early in development (Dong et al., 2009), and to “bootstrap” motion-dependent networks in the tectum. In older animals, nuanced motion-detecting networks could serve as a first line of defense, identifying early phases of looming, and allowing precise course corrections (Khakhalin et al., 2014; Bhattacharyya et al., 2017), while dimming detectors remain as a backup, mediating last-moment, less coordinated responses. Moreover, every time collision avoidance is not performed perfectly, sensorimotor networks can be refined, based on the mechanosensory inputs from the lateral line and from the skin (Felch et al., 2016; Helmbrecht et al., 2018).

In this study, we use high-speed calcium imaging and functional connectivity reconstruction to search for looming detectors within recurrent networks in the developing tectum of *Xenopus* tadpoles. Tadpole tectum is uniquely suitable for studies of sensory integration, as it is excessively plastic (Pratt and Aizenman, 2007; Busch and Khakhalin, 2019), strongly interconnected (James et al., 2015), and develops reliable looming selectivity within about a week (Dong et al., 2009; Khakhalin et al., 2014), as tadpoles mature from Nieuwkoop stage 45 to stage 49. We hypothesize that looming detectors may emerge in development, when connections between tectal cells are reshaped by patterned visual stimulation. Synapses in the tectum of young tadpoles exhibit spike-time-dependent plasticity (STDP; Zhang et al. 1998; Mu and Poo 2006; Vislay-Meltzer et al. 2006; Richards et al. 2010) that is expected to promote the development of so-called synfire chains: groups of neurons, sequentially connected to one another (Fiete et al., 2010; Zheng and Triesch, 2014). Synfire chains are selective for inputs that activate neurons in the same sequence in which they are connected (Clopath et al., 2010), turning them into temporal pattern detectors. We hypothesize that it is this ability to “resonate” with specific temporally patterned inputs that underlies looming behavior in late pre-metamorphic tadpoles.

There are two potential ways to test this hypothesis. First, we can look for synfire chains in the tectum; see whether they get selectively activated in response to looming stimuli, and check whether their connectivity and selectivity patterns change with development. Luckily, signal transmission in *Xenopus* tectal neurons is relatively slow, with synaptic transmission and spiking initiation taking up to 10 ms (Ciarleglio et al., 2015; Jang et al., 2016; Busch and Khakhalin, 2019), so if synfire chains in the tectum exist, they may be directly observable with fast calcium imaging, operating at rates of ∼100 frames/s. Second, if synfire chains can serve as a basis for looming detection, we can expect looming selectivity to spontaneously emerge in a model of a developing tectum governed by spike-time-dependent plasticity (Gao and Ganguli, 2015; Pietri et al., 2017). We can then use this model to explore the range of developmental rules that enable looming selectivity (Linderman and Gershman, 2017; Bassett et al., 2018), similar to recent advances in modeling the development of visual receptive fields (Bashivan et al., 2018), grid cells (Banino et al., 2018), and decision circuits (Haesemeyer et al., 2018).

With this in mind, we ask three specific questions. First, we test whether it is possible to reconstruct functional connectivity in tectal networks from calcium imaging recordings. Then we investigate the topology of reconstructed connectivity graphs, and compare these graphs between younger and older tadpoles. Finally, we check whether looming detection emerges in a computational model, and whether the statistics of simulated networks matches that of biological networks.

## Results

For all analyses, we name and interpret them in the text, but leave precise definitions for the Methods section. P-values are reported without correction, and are interpreted according to Fisher, rather than Neyman-Pearson philosophy (Greenland et al., 2016; Amrhein et al., 2019), with more attention payed to hypotheses supported by several alternative analyses. All code and summary data are available at: https://github.com/khakhalin/Ca-Imaging-and-Model-2018.

We performed calcium imaging experiments with cell-permeant Oregon Green Bapta in 14 stage 45-46, and 16 stage 48-49 tadpoles, recording responses from 128 ± 40 tectal cells per experiment (between 84 and 229; here and below “ ± “ after the mean denotes standard deviation). Unless stated otherwise, sample sizes *n* = 14 and 16 animals for stage 46 and 49 tadpoles apply to all analyses between younger and older animals in this study. To each tadpole, we presented a sequence of three different stimuli, always in the same order: a dark-on-light “Looming” stimulus, followed by a full-field dark “Flash”, followed by a spatially “Scrambled” looming stimulus (Figure 1A). For a looming stimulus, a dark circle on a light background took 1 s to grow to an angular size of ∼70° (Khakhalin et al., 2014). Scrambled stimuli were dynamically identical to looming, except that the visual field was split into a 7×7 grid of square tiles, and these tiles were randomly rearranged. In total, to every animal we presented 60 ±11 stimuli (20 ±4 stimuli of each of three types). We recorded high speed (84 frames/s) calcium imaging signals (Xu et al., 2011; Truszkowski et al., 2017) from one layer of “deep” principal tectal neurons in the tectum (Figure 1B,C); extracted fluorescence traces (Figure 1D,E), and inferred instantaneous spiking rate for each neuron within every frame (Figure 1F,G).

**Figure 1.**
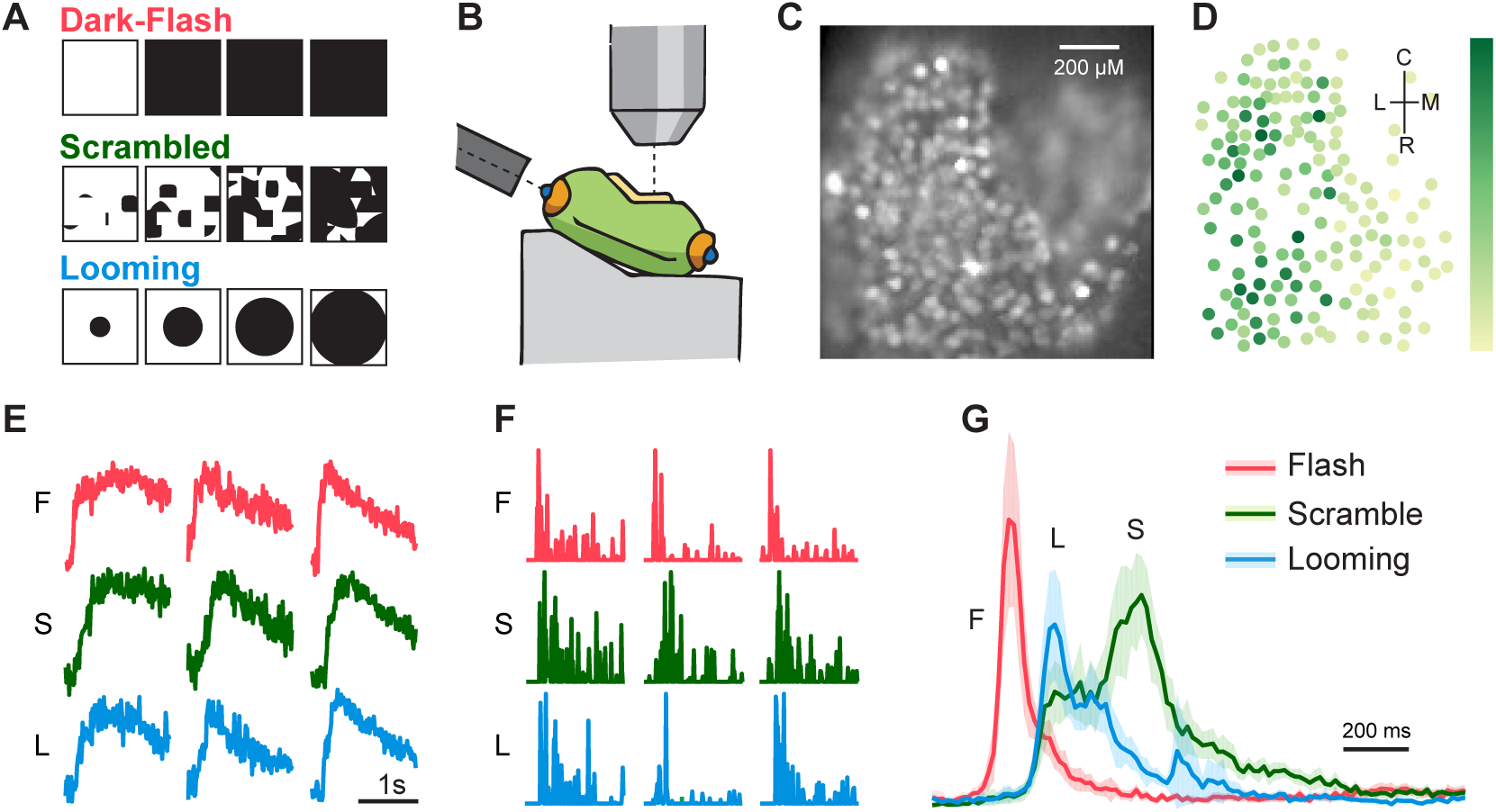
Experimental design overview. **A**. Visual stimulation: four representative frames from each stimulus type. **B**. Schematic of the preparation, with stimulation fiber on the left, and microscope objective on top. **C**. View of the optic tectum during calcium imaging recording. **D**. Regions of interest (cells) for brain shown in C, with darker markers representing cells with stronger responses. Labels in the right top corner mark Rostro-Caudal and Lateral-Medial axes. **E**. Typical fluorescence responses to Flash (F), Scrambled (S) and Looming (L) stimuli from three cells in the tectum. **F**. Spiking estimations (via deconvolution) for these fluorescence traces. **G**. Average full-brain responses to stimuli of every type, for one representative experiment, shown with a 95% confidence interval band.

### Responses and stimulus selectivity

As previously reported for electrophysiology experiments (Khakhalin et al., 2014), responses to flashes were fast, with a sharp peak, and little recurrent activation after the peak, while responses to looming stimuli were slower, and were followed by a strong recurrent activation (Figure 1G). Responses to both looming and scrambled stimuli were highly variable from one animal to another (Figure 2A), which may indicate either an inherent variability of network configurations, or different levels of inhibition across preparations.

**Figure 2.**
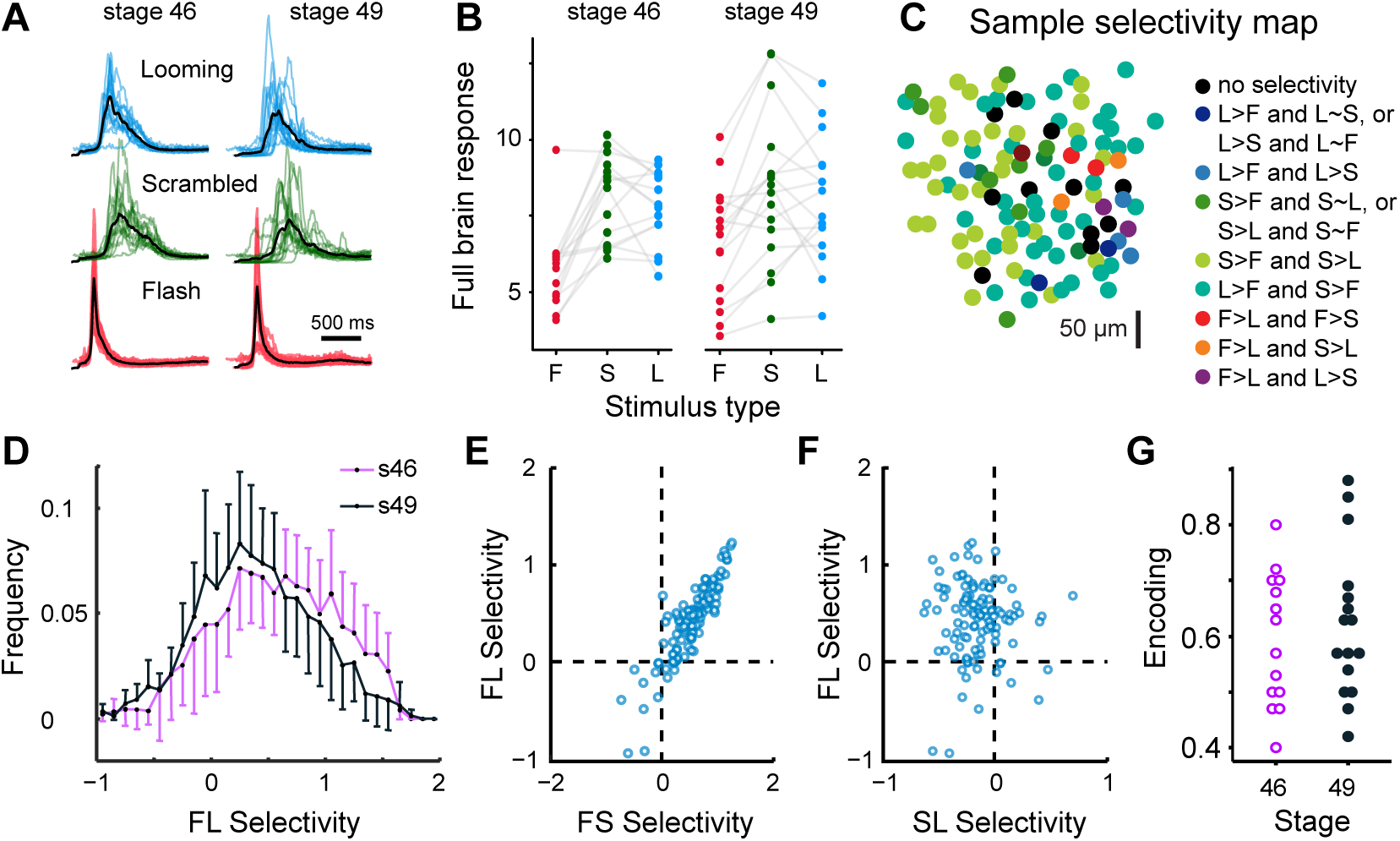
Selectivity analysis. **A**. Average brain responses (sum of activity of all recorded cells over time), for each stimulus type, in each experiment, superimposed. Black lines show grand averages across all experiments, separately for younger and older tadpoles. **B**. Cumulative full brain responses (an integral under the curves shown in A) for each experiment, across three stimulus types. **C**. A sample selectivity map, with hue coding preference for Flash (more red), Scrambled (more green), and Looming (more blue) stimuli. **D**. Average histograms of cell selectivity distributions, for younger (violet) and older (black) tadpoles; uncertainty bars show 95% confidence intervals, for clarity given in only one direction. Younger tadpoles had more highly selective cells than older tadpoles. **E**. A correlation between Flash-Scramble and Flash-Looming selectivity of individual cells in a typical experiment. **F**. A correlation between Scramble-Looming and Flash-Looming selectivity of individual sells in the same sample experiment. **G**. Stimulus encoding in different experiments, for two developmental stages.

The **total spiking output** of observed networks (the sum of all spikes of all neurons during a visual response) tended to be higher for looming stimuli than for flashes (Figure 2B; on average, 39±29% higher for younger; 25±25% higher for older tadpoles; *p*_*t*1_ = 2e-4 and 1e-3 respectively); this preference did not change in development (*p*_*t*_ = 0.15). There was no discernible difference in response amplitudes between looming and scrambled stimuli (average difference of −0.03±0.15 and −0.01±0.20; *p*_*t*1_ = 0.45 and 0.78 for younger and older animals respectively; no difference in development *p*_*t*_ = 0.80). These results match our prior reports that the overall strength of tectal responses in tadpoles depends mostly on the dynamics of visual stimuli, rather than on their geometry (Khakhalin et al., 2014; Jang et al., 2016).

Were there specialized looming detectors in the tectum? To quantify **stimulus selectivity** (Figure 2C), for each tectal cell we calculated Cohen’s effect size *d* between cumulative responses to different stimuli. We considered two types of selectivity: that for “Looming over Flash” (a type of selectivity that may rely on both stimulus dynamics and its spatial organization), and “Looming over Scrambled” (that can only rely on spatial properties of stimuli, as by design Looming and Scrambled stimuli had the same dynamics). Looking at cumulative responses was of course a simplification, as in real life animals respond to stimuli while they are still unrolling (Peron and Gabbiani, 2009; Khakhalin et al., 2014), and not after a timed 1 second-long presentation. We however had no way to justify a specific way of dynamic thresholding, and so opted for the simplest approach.

On average, tectal cells were selective for looming stimuli (Figure 2D; mean Cohen’s 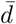 of 0.67 ±0.50 and 0.46 ±0.47 in younger and older tadpoles respectively). Here we first calculated selectivity *d* for every cell, then found average 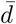 within each brain. There was no difference between stages in terms of either per-experiment mean selectivity 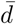, nor within-experiment variances in selectivity (*p*_*t*_ = 0.3, 0.3). The share of cells that responded to looming stimuli stronger than to flashes also did not change in development (84 ±23%, 77 ±21%; *p*_*t*_ = 0.4). Compared to younger brains, older brains had fewer highly selective cells (Figure 2D, right side of the curve). The gap between top-selective (90th percentile) and median selective cells was larger in younger (0.75 ±0.26) than in older tadpoles (0.53 ±0.27; *p*_*t*_ = 0.03), as selectivity distributions in younger tadpoles were more positively skewed (*p*_*t*_ = 0.01). These results were unexpected, as older tadpoles perform better in collision avoidance tests (Dong et al., 2009), and so we expected them to develop a subset of specialized looming-selective cells, as described in adult frogs (Nakagawa and Hongjian, 2010; Baranauskas et al., 2012), and other vertebrates (Wang and Frost, 1992; Wu et al., 2005; Liu et al., 2011). Yet in our experiments, a subpopulation of strongly selective cells not only did not expand in older animals, but became less prominent, somewhat similar to developmental changes described in the visual cortex (Rochefort et al., 2009).

We then considered a second, more computationally demanding definition of selectivity: a preference for spatially organized looming stimuli over scrambled stimuli. On average, tectal cells did not show a preference between these two stimuli (average selectivity of −0.07 ±0.33 in younger tadpoles, −0.04 ±0.49 in older ones; no change in development *p*_*t*_ = 0.9). The share of cells that responded to looming stronger than to scrambled was at a chance level for both developmental stages (46 ±31%, 48±37%, *p*_*t*_ = 0.9), and there was no change in either within-brain variance of this selectivity (*p*_*t*_ = 0.9), or the 90−50 percentile asymmetry of values (*p*_*t*_ = 0.8).

We found that selectivity for scrambled stimuli over flashes correlated with selectivity for looming stimuli over flashes (Figure 2E) in both developmental groups: average within-brain correlation coefficients *r* = 0.82 ±0.13, *p*_1*t*_ =3e-12 for younger animals, and 0.75 ±0.18, *p*_*t*1_ = 3e-11 older ones, with no change in development (*p*_*t*_ = 0.3). In contrast to that, the preference for looming over flashes did not correlate with preference for looming over scrambled (Figure 2F; *r* = 0.03 ±0.29, *p*_1*t*_ = 0.7 for stage 46; 0.13±.30, *p*_1*t*_ = 0.1 for stage 49). This further suggests that the majority of cells in the tectum responded to stimulus dynamics only, and did not specifically process its geometry.

Finally, as a holistic way to quantify tectal network selectivity, we looked at our ability to predict stimulus identity from all recorded responses in tectal cells (Avitan et al., 2016): a measure known as “**stimulus encoding**”. We ran a multivariate logistic regression on first half of the data, linking values of total responses of each cell in each trial to the type of stimulus used in this trial. Then we measured the quality of this linkage on the second half of recorded data (Figure 2G). We found that with the dynamics of responses disregarded (with predictions based only on the total spiking output of each cell), the quality of these prediction was rather low: 59 ±12% for younger, and 62 ±13% for older tadpoles, with no change in development (*p*_*t*_ = 0.6). ±

To assess the **variability of response shapes** from one tectal cell to another within each brain, we performed exploratory factor analysis (principal component analysis followed by a promax rotation) of responses to looming stimuli within each preparation. The first and second principal components explained on average 19 ±7% and 4 ±1% of variance in younger tadpoles, and 24 ±14% and 3 ±1% in older tadpoles, with second component encoding response timing (Figure 3A). Cells with early responses to looming stimuli were reliably grouped together somewhere within the recorded field (Figure 3B) in each one out of 30 experiments (see Methods). We assumed that early-responding cells grouped together because of the known retinotopic arrangement of receptive fields in the tectum (Ruthazer and Cline, 2004), that made tectal activation follow and reproduce the gradual unrolling of looming stimuli on the retina. Across cells, the latency of average response correlated with distance from the retinotopy center (Figure 3C; *r* = 0.35 ±0.24; correlations individually *p*_*r*_ < 0.05 in 25/30 experiments), despite our latency estimations being quite noisy for low-amplitude cells (see Methods). Curiously, while visual projections to the tectum are known to be actively refined in development (Sakaguchi and Murphey, 1985; Ruthazer and Cline, 2004; Munz et al., 2014), it seems that the precision of functional retinotopic maps did not differ between younger and older tadpoles, as the values of correlation coefficients between cell position and early component prominence did not discernibly change in development (*r* = 0.63 ±0.21 and 0.57 ±0.25 respectively, *p*_*t*_ = 0.5), which is similar to prior reports in Zebrafish (Avitan et al., 2016).

**Figure 3.**
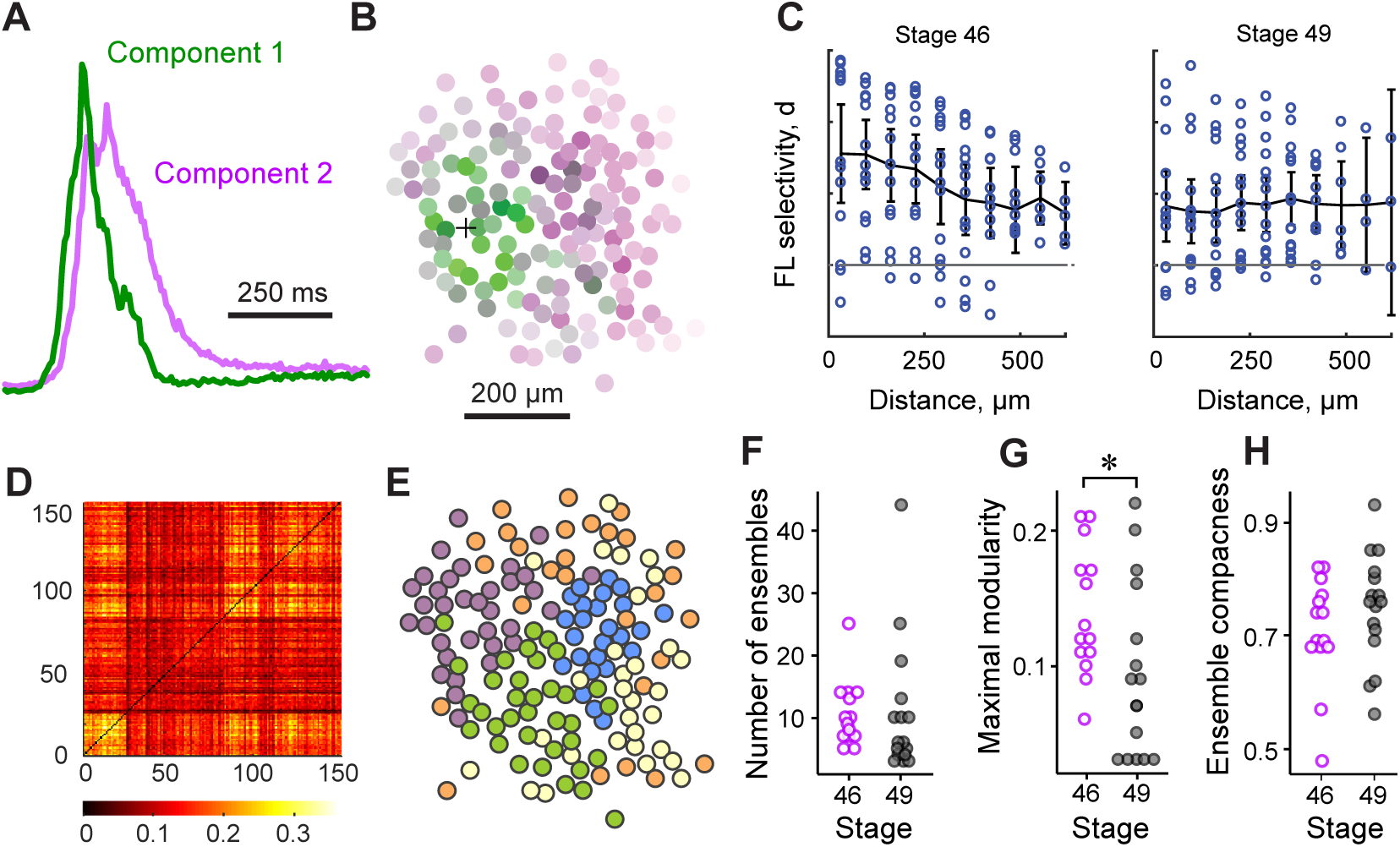
Spatiotemporal organization of responses. **A**. Two first components (Principal Component Analysis with promax rotation) identified in tectal responses in a typical experiment. **B**. Neurons from the same sample experiment as in A, shown as they were located in the tectum, and colored by the contribution of early (green) and late (purple) response components. Black cross shows the estimated position of the retinotopy center. **C**. Average change of Flash-Looming selectivity with distance from the retinotopy center, for stage 46 (left) and stage 49 (right) tadpoles; uncertainty bars show 95% confidence intervals. **D**. Adjusted correlation matrix for instantaneous activation of different neurons across all stimuli in a sample experiment (same as in B). **E**. Ensembles identified from the correlation matrix. **F**. The number of ensembles, identified in younger and older tadpoles. **G**. Maximal modularity of ensemble partition. **H**. Ensemble spatial compactness.

Knowing where the center of a looming stimulus was projected within the tectum, we could check whether looming-selective cells tended to be found in the center of the expanding activation area (as it would be expected if looming detectors formed a meta-retinotopic map; Frost and Sun 2004), or at the periphery. We found that selectivity for looming over flash tended to decrease with distance from the projection center (Figure 3C) for both stage 45 (average *r* = −0.37 ±0.27; individual correlations *p*_*r*_ <0.05 in 12/14 experiments), and stage 49 tadpoles (average *r* = −0.09 ±0.35; *p*_*r*_ <0.05 in 12/16 experiments), indicating that looming-selective were often found in the center of the emerging spatial response. Similarly, at both developmental stages, selectivity decreased with response latency (stage 46: *p*_*r*_ <0.05 in 11/14 animals, average *r* = −0.29 ±0.11; stage 49: *p*_*r*_ <0.05 in 10/16 animals, average *r* = −0.16 ±0.21). Both correlations were weaker in older tadpoles (*p*_*t*_ = 0.02 for a link between distance and selectivity, *p*_*t*_ = 0.03 for a link between response latency and selectivity), suggesting a more spatially even distribution of looming-selective cells in older tectal network.

### Neuronal ensembles

We then asked whether certain tectal cells tended to respond or stay silent together, potentially indicating distributed processing in network ensembles (Orger, 2016). As a simplistic way to assess that, we performed factor analysis of trial-to-trial **population response variability**, and checked whether its results changed in development, as it happens for tectal spontaneous activity (Xu et al., 2011). The total number of principle components needed to describe 80% of variance across cells (Avitan et al., 2017) was similar in stage 46 and 49 tadpoles (51±14 and 65±28, *p*_*t*_=0.1 for looming; 49±12 and 62±32, *p*_*t*_=0.2 for flashes; 51±14 and 64±26, *p*_*t*_=0.1 for scrambled).

To better describe and visualize network activation variability, we looked for **ensembles** of tectal cells using spectral clustering (Thompson et al., 2016). Unlike for spontaneous activity, we could not easily aggregate activity states into clusters (Avitan et al., 2017), as activation in our networks was driven by shared sensory inputs. Instead, we subtracted average responses of each cell from its responses in individual trials, and calculated pairwise correlations on the remaining “anomalies” of trial-by-trial activation (Figure 3D). We turned these correlations into pairwise distances in a multidimensional space, ran a series of spectral clustering partitions (Ng et al., 2002), and of all possible partitions, picked the one that maximized spectral modularity on a similarity graph (Newman, 2006; Gómez et al., 2009) (Figure 3E; see Methods for details). We found that the number of ensembles did not differ between younger and older tadpoles (Figure 3F; 10 ±5 in stage 45, 11 ±11 in stage 49; *p*_*t*_=0.9), although in older tadpoles, ensembles were more coordinated with each other, producing lower values of network modularity (Figure 3G; 0.14 ±0.05 to 0.09 ±0.06, *p*_*t*_=0.03). Tectal ensembles also tended to be spatially localized (Figure 3H), with cells within each ensemble located closer to each other on average than to cells from different ensembles, both in younger (29 ±9% closer) and older animals (25 ±10% closer; no difference in development *p*_*t*_=0.3).

### Network reconstruction

The high speed of imaging used in this study (84 frames/s) allowed us to look not just at instantaneous correlations between activation of individual neurons, but at the *propagation* of signals through the network. To reconstruct network connectivity, we calculated pairwise transfer entropy (Gourévitch and Eggermont, 2007; Stetter et al., 2012) between activity traces of individual cells. Intuitively, for each pair of neurons *i* and *j* we quantified the amount of additional information that past activity of neuron *i* could offer to predict current activity of neuron *j*. This is conceptually similar to calculating a cross-correlation between the activity of neuron *i* at each frame *t*, and the activity of neuron *j* at each consecutive frame *t* + 1 (Figure 4A), except that transfer entropy calculation (Figure 4B) does not make assumptions about the type of influences neuron *i* may have over neuron *j*, and offers a lower inference noise (Stetter et al., 2012).

**Figure 4.**
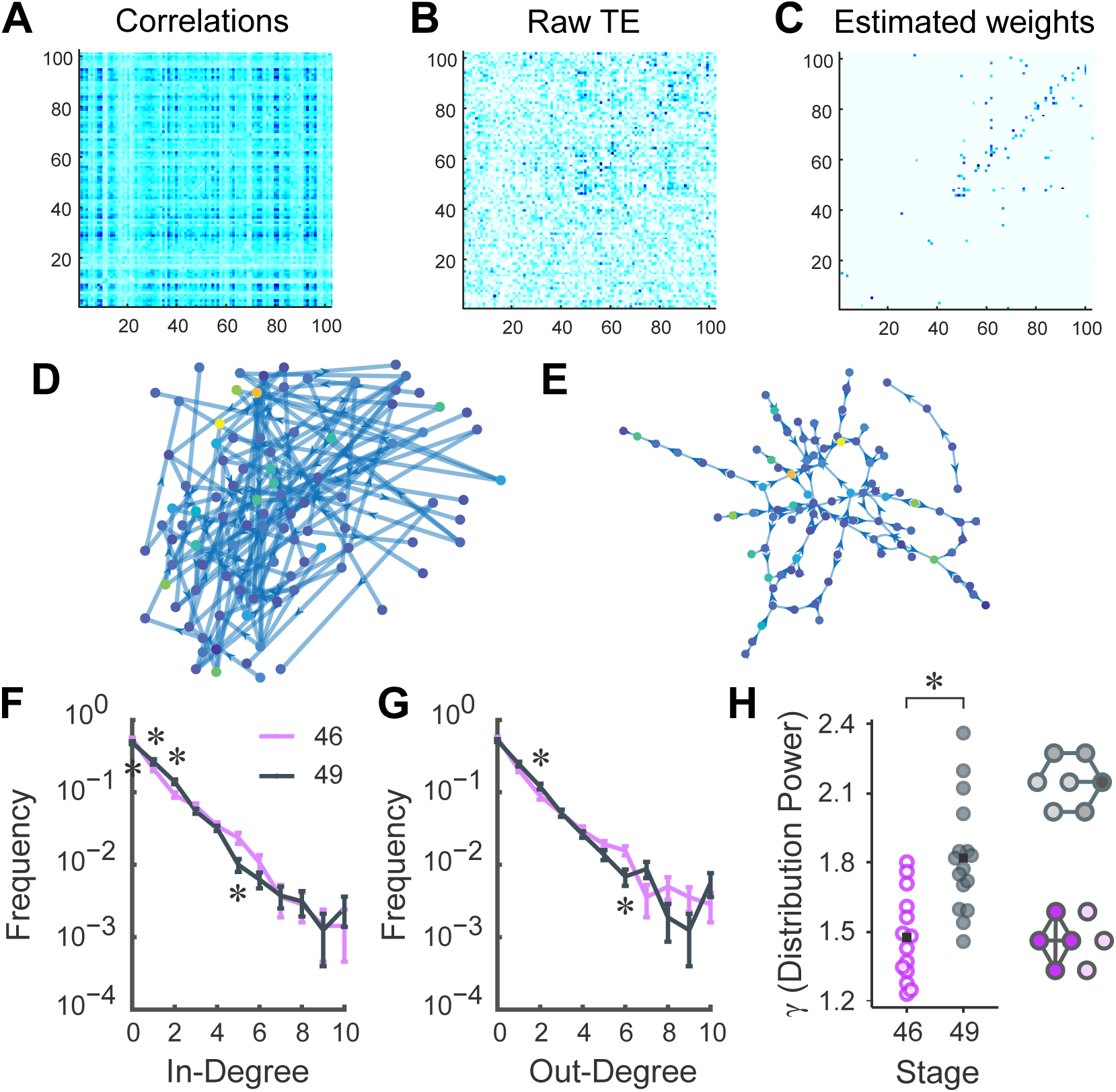
Connectivity reconstruction. **A**. Delayed correlations between activities of different neurons in a sample experiment. **B**. Adjusted transfer entropy estimations for the same experiment. **C**. Estimated weighted adjacency matrix. **D**. Connectivity reconstruction, with location of each cell preserved. Node color indicates its selectivity. **E**. Same graph as in D, in visually optimized layout. Positions of individual nodes no longer represent the position of tectal cells. **F**. In-degree distribution for younger (violet) and older (black) tadpoles, presented in log-scale, with uncertainty bars showing 95% confidence intervals. Observable changes for individual degrees (*p*_*t*_ < 0.05) are marked with asterisks. **G**. Same as C, but for out-degree distributions. **H**. Network power *γ* increased in development, as older tadpoles had a slightly sharper slope to their degree distributions.

In our experiments, all tectal neurons received shared inputs from the eye, that recruited them in a similar manner in every trial. This made the connectivity inference more complicated, as neurons could spike in a sequence both because they were connected, and because they received innervation from sequentially activated areas of the retina (Mehler and Kording, 2018). To compensate for shared inputs, we randomly reshuffled trials for every neuron, calculated average transfer entropy on reshuffled data, and subtracted it from the value obtained on trial-matched data (Gourévitch and Eggermont, 2007; Wollstadt et al., 2014). In essence, it is similar to working with deviations from the average response, and quantifying whether these deviations tended to propagate through the network, from one neuron to another. The reshuffling step also allowed us to calculate *p*-values for each pair of neurons, and quantify how unusual the value of transfer entropy was, compared to a value arising from shared inputs alone.

We interpreted transfer entropy values as approximations of weights in a connectivity matrix **W**, with *w*_*ji*_ describing the strength of connection from neuron *i* to neuron *j* (but see Mehler and Kording 2018). We calculated **W** and corresponding *p*-values independently on looming, flash, and scrambled stimuli, and used these independent estimations to ensure some level of internal replication within each experiment (see Methods). Of all possible edges, only 1.6 ±1.4% were found to be non-zero in all three independent analyses, which was discernibly higher than 0.6 ±0.09% expected if edges were assigned at random (paired t-test *p*_*tp*_ = 9e-9). In each experiment, we then introduced cut-offs on edge *p*-values, to exclude weak noisy edges from the connectivity graph (Figure 4C). As noise levels varied across preparations, we adjusted these cut-offs (Stetter et al., 2012), making the number of non-zero edges in each reconstruction equal to the number of recorded cells (Figure 4D,E). The effective *p*-value cut-offs were between 0.001 and 0.007 depending on the experiment (median of 0.004). Note also that the results described below are relatively insensitive to the precise assumptions of this approach to weigh thresholding.

The simplest statistical property of a network is its **degree distribution**: the share of nodes with different numbers of incoming (*k*_*in*_) and outgoing (*k*_*out*_) connections. We compared rounded degree distributions between networks detected in younger and older tadpoles (Figure 4F,G), and found that older networks contained fewer unconnected cells (*k*_*in*_ = 0, *p*_*t*_ <0.03) and fewer cells with high number of connections (*k*_*in*_ = 5, *k*_*out*_ = 6, *p*_*t*_ = 0.01 in both cases), but more cells with intermediate number of connections (*k*_*in*_ = 2, *p*_*t*_ = 0.001, and *k*_*out*_ = 2, *p*_*t*_ = 0.04). When we approximated degree distributions (excluding *k* = 0) with a power law *P* (*k*) ∼ *k*^*−γ*^, the power constant *γ* was smaller in younger (1.48±0.19) than in older tadpoles (1.82±0.25, *p*_*t*_ = 2e-4; Figure 4H), consistent with a steeper drop from the rate of occurrence of weakly connected to highly connected cells. It suggests that older tectal networks had more chains of connected neurons (out- or in-degree of *k* = 1) and forks (*k* = 2), while younger neurons had more hyperconnected hubs (*k* >5) and unconnected nodes (*k* = 0), which matches expectations for STDP-driven networks (Fiete et al., 2010).

An unusual feature of our high-speed imaging protocol, compared to more common techniques that rely on genetically encoded calcium sensors and confocal microscopy, was that the signal-to-noise ratio varied greatly from one cell to another, depending on how far the cell was from the focal plane, and how much dye it absorbed through the partially exposed membrane during staining. As a result, the share of cells with weak signals that appeared unconnected to the rest of the network varied across experiments, and was dependent on extraneous circumstances, such as the physical curvature of the preparation. To ensure that poorly resolved cells did not bias our conclusions too strongly, we restricted further network analysis to the largest weakly connected component of each network. There was no difference in the number of weakly connected components detected in younger and older tadpoles (50 ±14 and 50 ±26 for stages 45 and 49 respectively, *p*_*t*_ = 0.9), but in older animals the largest weakly connected component included a slightly higher share of observed cells (50±6% and 64±12% for younger and older networks respectively; *p*_*t*_ = 4e-4), consistent with the change in degree distributions described above.

A known prediction for networks dominated by spike-time-dependent plasticity (STDP) is that with time, neuronal connections tend to become highly asymmetric (Pratt et al., 2008; Richards et al., 2010). Indeed, if cells *i* and *j* are reciprocally connected, every time *j* spikes after *i*, STDP would increase the weight *w*_*ji*_ (for a connection leading from *i* to *j*), but decrease the reciprocal weight *w*_*ij*_ (Abbott and Blum, 1996; Fiete et al., 2010). We found that for our data, **the share of bidirectional edges** (those with both *w*_*ij*_ and *w*_*ji*_ >0) among all detected edges was smaller (0.3±0.3%) than expected for random edge assignment in graphs of our size (0.4±0.1%, paired *p*_*t*_ = 0.02), indicating an asymmetric information flow in the tectum. Moreover, the share of bidirectional edges decreased in development, from 0.4±0.3% in younger animals to 0.2±0.2% in older animals (*p*_*t*_ = 0.03), suggesting that STDP was still shaping emerging network topology at these developmental stages.

We then looked at whether connected cells were more likely to be located spatially closer to each other. We found that the **average distance between connected cells** was indeed shorter than for randomly selected cells (18±10% shorter for stage 45, and 17±8% shorter for stage 49 tadpoles; connected neurons were closer to each other with *p*_*t*_ <0.05 in 13/14 and 16/16 individual experiments respectively). In contrast to visual inputs to the tectum (Tao and Poo, 2005), the intra-tectal connectivity did not become more compact in development (*p*_*t*_=0.7). This may suggest that tectal networks rely on relatively far-reaching recurrent connections to integrate visual information across the visual field (Baginskas and Kuras, 2009; Liu et al., 2016; Jang et al., 2016).

### Network properties

The best (and perhaps the only possible) way to compare two sets of irregular connectivity graphs to each other is to quantify their properties using a diverse set of network measures, and compare these values across groups. We did this, while also checking whether our inferred connectivity graphs were statistically unusual, by comparing their properties to that of matching randomized graphs (Ansmann and Lehnertz, 2012). To do so, we randomly rewired edges between nodes, while keeping the distribution of edge weights *w*_*ji*_, and the number of non-zero edges leaving each node (node out-degree) fixed, as a case of degree-preserving rewiring (Maslov and Sneppen, 2002) generalized for weighted directed graphs.

The measure of average “connectedness” in the graph, known as **network efficiency**, quantifies the length of the average shortest path connecting two random nodes in the network (Latora and Marchiori, 2001; Rubinov and Sporns, 2010). This value is high when short paths made of high-weight edges tend to connect any two randomly chosen nodes in the graph, making it easy for signals to propagate within the network; the value is low when some pairs of nodes are far from each other on a graph (Figure 5A). We found that network efficiency (0.004±0.002 for stage 46, 0.002±0.002 for stage 49 tadpoles) was lower than expected for a randomized network with a matching degree distribution (*d* = −0.3, paired *p*_*t*_ = 0.04 and *d* = −0.3, paired *p*_*t*_ = 0.06 for younger and older tadpoles respectively). It was not clear whether efficiency changed with age (*d* = −0.8, *p*_*t*_ = 0.06).

**Figure 5.**
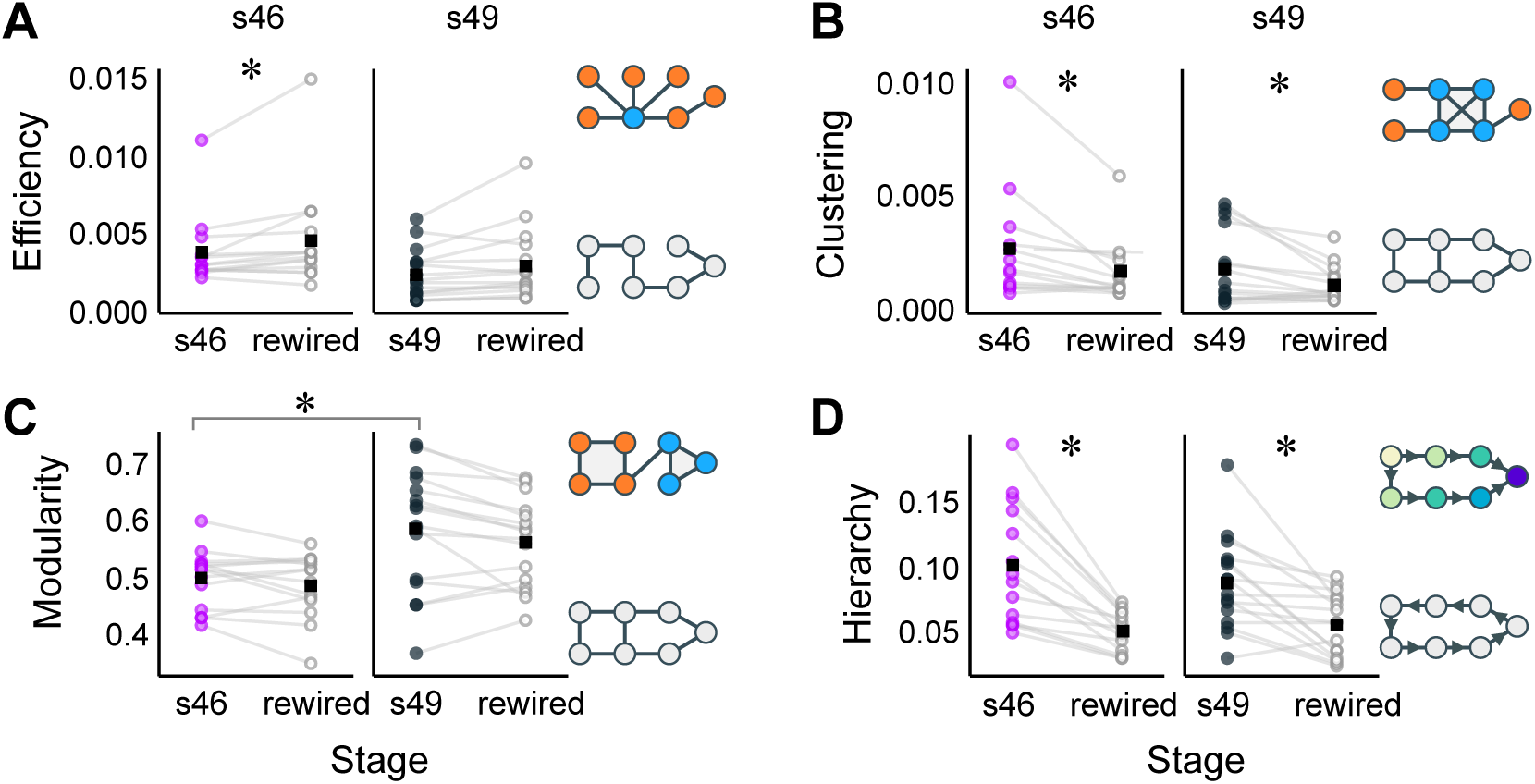
Global network properties. Filled markers show values from experiments; empty markers show averaged values for matching rewired networks; black squares show averages for each group. Network diagrams to the right of each plot show cartoon representation of graphs with low and high values of respective network measures. Asterisks show statistically discernible differences (*p* < 0.05). **A**. Global network efficiency was lower than expected for stage 46 tadpoles. **B**. Global clustering was higher than expected by chance, for both developmental stages. **C**. Network modularity increased in development. **D**. Network hierarchy was higher than expected by chance, for both developmental stages.

The **global clustering coefficient** describes the small-scale heterogeneity in the network (Fagiolo, 2007), and is defined as the relative frequency of two neighboring nodes being a part a triangle with a third node connected to both of them (Figure 5B). The value of clustering coefficient in our networks was very small (2.4±2.5 e-3 for stage 46, 1.5±1.6 e-3 for stage 49 animals), but slightly larger than expected in a randomly rewired network with same degree distribution (*d* = 0.5 and 0.6, paired *p*_*t*_ = 0.01 and 0.02 for younger and older animals). This means that neurons with more than two connections were likely to form clusters. There was no change in clustering in development (*d*=−0.4, *p*_*t*_=0.3, Fig.).

Network **modularity** is the most commonly used measure of mesoscale network heterogeneity (Newman, 2006; Leicht and Newman, 2008). A network with high modularity can be split into a set of sub-networks, with higher density of connections within each sub-network, and weaker connections between them (Figure 5C). In our experiments, reconstructed tectal networks had similar, or slightly higher modularity, compared to matching randomly rewired networks (*d* = 0.2 and 0.3, paired *p*_*t*_ = 0.2 and 0.06, for stages 46 and 49 respectively), and network modularity clearly increased in development (*d* = 1.0, *p*_*t*_ = 0.01).

#### Flow hierarchy

is a measure of structural hierarchy in the network (Mones et al., 2012), assessed through flows of activation that propagate through it. We based our measure of hierarchy on the distribution of Katz centrality values (Katz 1953; Fletcher and Wennekers 2018; see next section for definitions). Intuitively, hierarchy is high when a network has groups of “input” and “output” nodes, with connections between them largely pointing in the same direction, as it happens in layered feed-forward networks; hierarchy is weak in random networks, or networks that consist of cycles with no clear inputs and outputs (Czégel and Palla, 2015). We hypothesized that a network of dedicated looming detectors may exhibit flow hierarchy, with more edges leading from “feeder neurons” to “detector neurons”. Indeed, tectal networks were more hierarchical than randomized networks with matching degree distributions (Figure 5D; *d* = 1.5 and 1.1, paired *p*_*t*_ = 1e-04 and 1e-03 for younger and older tadpoles respectively). There was no difference in flow hierarchy in development (*d* = −0.3, *p*_*t*_ = 0.4).

### Selectivity Mechanisms

Even though the exact architecture of collision-detecting tectal networks is unknown, it is safe to assume that looming-selective neurons have to integrate streams of information coming from different parts of the visual field. We therefore hypothesized that the topological placement of selective neurons within each connectivity graph would not be random (Timme et al., 2016). To verify that, for each reconstructed network we tested correlations between looming selectivity of each neuron and values that quantify its placement within the network, known as measures of **node centrality**.

To identify information sinks (neurons that tended to collect activation from the rest of the network), for each cell we calculated its **Katz centrality** within the graph (Katz, 1953; Fletcher and Wennekers, 2018). By definition, nodes with high Katz centrality have many paths leading to them, so a spike originating at random within a graph is more likely to eventually cause activation of these nodes, compared to nodes with low Katz centrality. We found that when all cells from all experiments were considered (Figure 6A), there was a very weak but statistically discernible correlation between the looming selectivity of cells and their Katz centrality (*r* = 0.02, *p*_*r*_ = 1e-6, *n* =2487). More convincingly, correlation coefficients between Katz centrality and looming selectivity were weak but positive in 19/30 experiments (Figure 6B; average *r* = 0.09±0.20 *p*_*t*1_ = 0.03). There was no change of this effect in development (*p*_*t*_ = 0.5).

**Figure 6.**
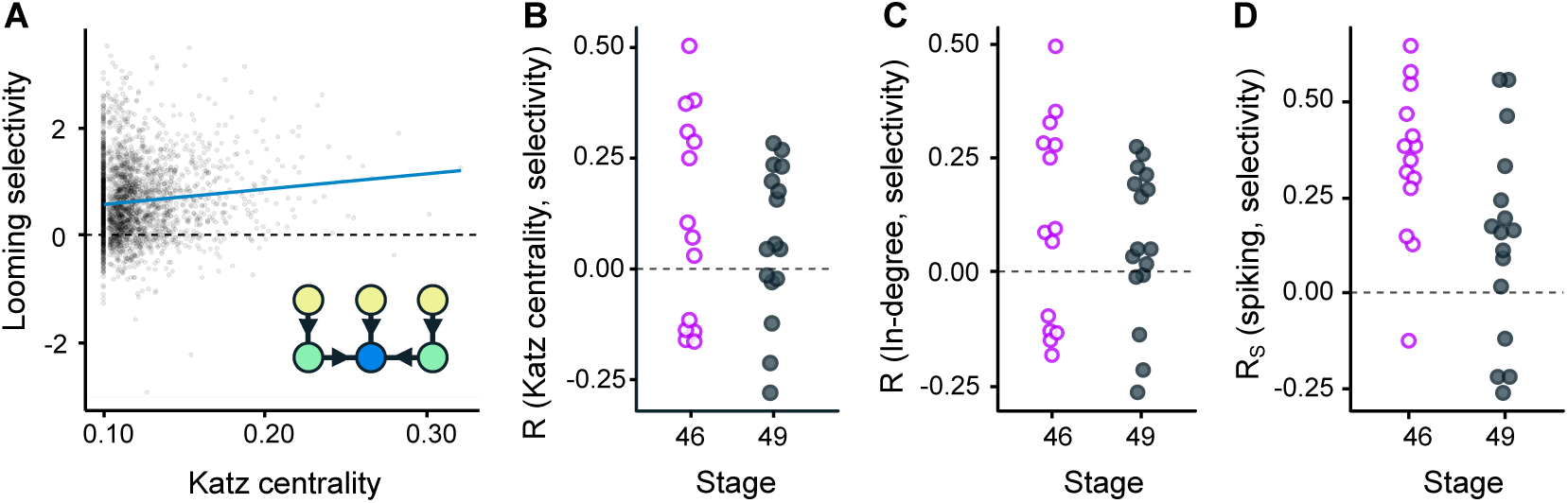
Local network properties (centrality) for cells selective for looming stimuli. **A**. Cells with higher Katz centrality had a weak tendency to respond stronger to looming stimuli (*r* = 0.02, *p*_*r*_ = 1e-6). A diagram inset illustrates the concept of Katz centrality: cells with low centrality (sources) are pale, while cells with higher centrality (sinks) are darker. **B**. Correlation coefficients for Katz centrality of each cell and its selectivity for looming stimuli, calculated for each experiment, and shown by developmental stage. **C**. Correlation coefficients for node in-degree and its selectivity for looming stimuli, in each experiment, shown by developmental stage. **D**. Correlation coefficients for average activation of each cell, and its selectivity for looming stimuli, in each experiment, shown by stage.

A node may have high Katz centrality for two reasons: it may receive a higher number of incoming connections (higher **in-degree**), or have chains of directed edges leading to it. To see which of these patterns may be at work here, we checked whether looming-selective cells tended to receive more incoming connections, compared to non-selective cells (Litwin-Kumar and Doiron, 2014). When all points were considered, there was no correlation between in-degree and cell selectivity (*p*_*r*_ =0.3, *n* = 2487), but for individual experiments, correlation coefficients between in-degree and selectivity were positive in 20/30 of cases (Figure 6C, *p*_*t*1_ = 0.03), which suggests that selective cells may tend to receive more incoming connections than non-selective cells. There was no change of this correlation in development (*p*_*t*_ = 0.5).

As cells with higher Katz centrality tend to be activated more often (Fletcher and Wennekers, 2018), we checked whether selectivity for looming stimuli correlated with cell **average spiking activity** in our recordings. We found that both in younger and older animals, actively spiking cells tended to have stronger looming selectivity (for stage 46 average *r*_*S*_ = 0.34±0.20, *r* > 0 with *p*_*t*1_ = 2e-5; for stage 49 *r*_*S*_ = 0.14±0.26, *p*_*t*1_ = 0.04, indicating a discernible decrease with development *p*_*t*_ = 4e-4; across all cells *r*_*S*_ = 0.07, *p*_*r*_ = 4e-4; Figure 6D). This result also matches the predictions from our studies of intrinsic excitability in the tectum (Busch and Khakhalin, 2019), where we show that higher spiking cells tend to be selective for slower synaptic inputs. For the analysis above we used Spearman rather than Pearson correlation, as 4 experiments had single neurons (one per experiment) with extremely high activity levels that acted as influential points. Excluding these 4 neurons out of 2487 total and using Pearson correlation yielded similar results.

If looming-selective cells gather information from the network, it was plausible to expect that they could be connected to other selective cells more often than to non-selective ones. To test this, we looked into **assortativity of selectivity**: a weighted correlation between selectivity scores of cells connected by edges, with strength of these edges taken as weights (see Methods). We found that in both younger and older tadpoles, selectivity values for connected cells correlated (for stage 46: *r* = 0.07±0.13, *p*_*t*1_ = 0.04, individual *r* > 0 in 9/14 experiments; for stage 49: *r* = 0.07±0.10, *p*_*t*1_ = 0.02; individual *r* > 0 in 10/16 experiments). This suggests that similarly selective cells indeed tended to be connected to each other. There was no change in this effect over development (*p*_*t*_ = 0.9).

Finally, we checked whether higher average activity of looming-selective cells linked them into tight clusters, or “rich clubs”. We calculated **local clustering coefficient** for each cell, and checked whether it correlated with cell selectivity. We found that local clustering coefficient did not correlate with cell selectivity both when all cells were pooled together (*p*_*r*_ = 0.6, n=2487), or across individual reconstructions (correlation coefficients across experiments were not different from zero; *p*_*t*1_ = 0.25, n=30). This shows that while selective cells tended to be connected to each other, they did not form clusters. Together these results suggest that the distribution of selective cells in tectal networks was not random.

### Developmental Model

To provide a counterpoint to our experimental results, we built a mathematical model of the developing tectum, and ran simulated responses from this model through the *exactly same* set of analyses that we used for biological experiments. The model consisted of 81 artificial neurons, arranged in a 9×9 grid (Figure 7A), that were all originally connected to each other (every neuron to every neuron) with random positive (excitatory) synaptic weights (Figure 7B). The model operated in discrete time, in 10 ms increments, and we interpreted the output of each neuron at each time step as its instantaneous firing rate. At each time step we looked at the network activity at the previous step, calculated total synaptic inputs to each neuron, and used a sliding logistic function to translate these synaptic inputs to postsynaptic spiking (see Methods for a full detailed description).

**Figure 7.**
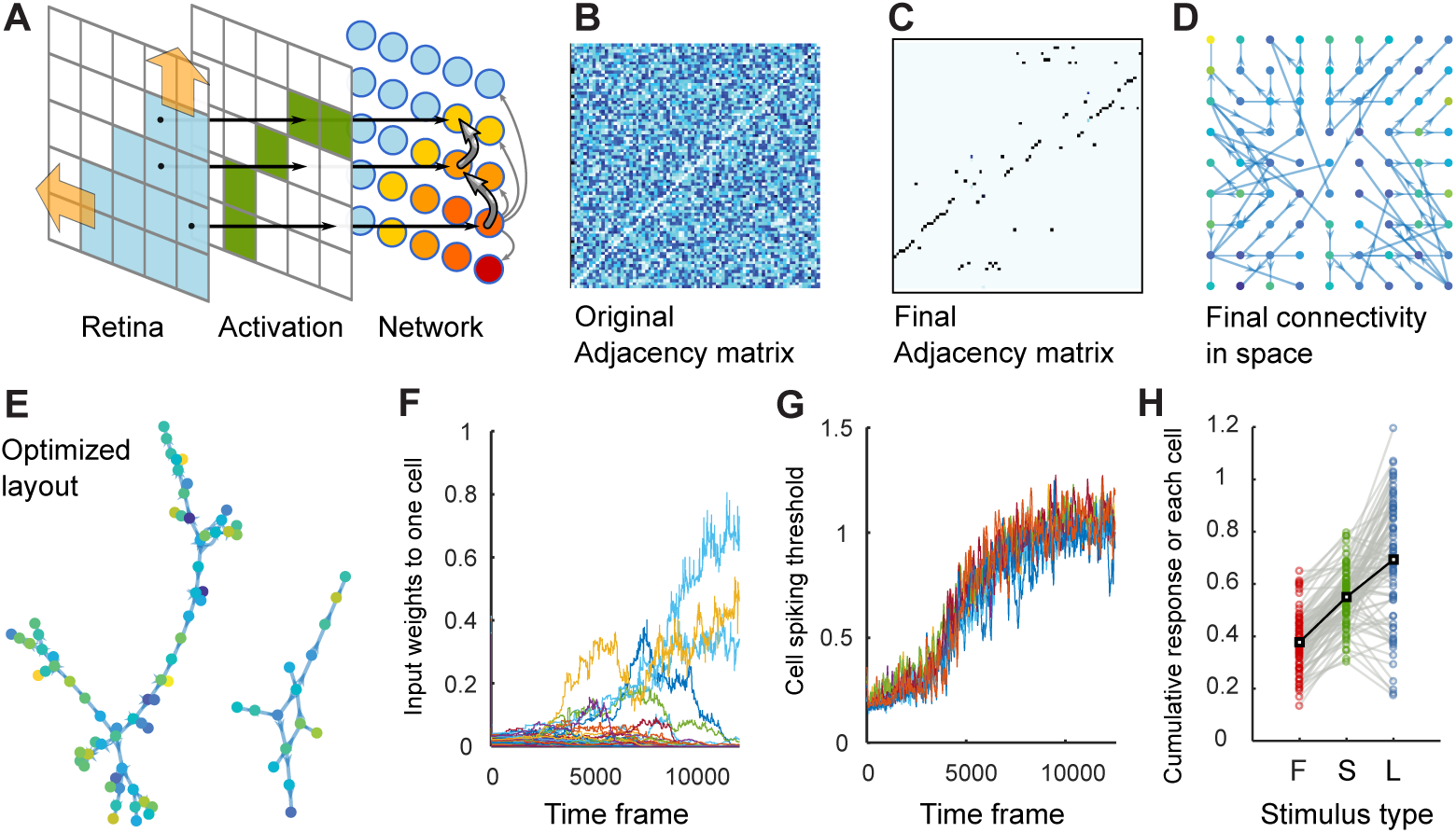
Developmental model. **A**. Model diagram (see the text). **B**. Typical adjacency matrix before simulation (random seed). **C**. Typical adjacency matrix by the end of simulation. **D**. Visualization of neuronal connections in space, with cell positions properly represented. Green and yellow nodes are selective for looming stimuli. **E**. Same graph as in C, but rearranged to reveal its structure. **F**. Evolution of all input weights to one sample cell, over the course of simulation. **G**. Evolution of spiking threshold for 9 sample cells, over the course of simulation. **H**. Cumulative responses of every cell to Flash, Scrambled, and Looming stimuli in a sample simulation.

We introduced three simple developmental rules: spike-time-dependent plasticity (STDP), homeostatic plasticity, and synaptic competition. Our implementation of STDP approximated biological STDP observed in the tectum (Zhang et al., 1998; Mu and Poo, 2006): if two cells were active one after another in two consecutive time frames, and they were connected with a synapse, the weight of this synapse was increased. Conversely, if two cells were active within the same time frame, the weights of a synapses connecting them were decreased, as it means that cells would try to activate each other immediately *after* they spiked (see Figure 7F for a typical evolution of synaptic inputs to one sample cell). The homeostatic plasticity rule adjusted excitability thresholds, trying to keep spiking output of each neuron constant on average (Pratt and Aizenman 2007; Turrigiano 2011; Figure 7G). Synaptic competition attempted to keep close to a constant both the sum of synaptic inputs to each neuron, and the sum of outputs from each neuron. In practice, it means that every time a synapse connecting neurons *i* and *j* increased in strength, all output synapses of neuron *i* and all input synapses of neuron *j* were scaled down a bit (Cohen-Cory, 2002; Munz et al., 2014; Hamodi et al., 2016).

With these three rules at play, we exposed the model to patterned sensory stimulation that imitated retinotopic inputs from the eye (Figure 7A). We hypothesized that during collision avoidance in real tadpoles, STDP-driven changes in the tectum may be controlled and amplified by a global learning signal that arrives if collision avoidance was unsuccessful (Savin and Triesch, 2014; Aswolinskiy and Pipa, 2015; Heap et al., 2018b), and originates either in dimming receptors in the retina (Baranauskas et al., 2012), or in mechanosensory systems of the hindbrain (Pratt and Aizenman, 2009; Felch et al., 2016; Truszkowski et al., 2017). This approach is known as the eligibility trace model, in which changes in synaptic weights do not happen immediately, but are first “remembered” by each cell as potential changes, and are only implemented in response to a timely reinforcement signal (Seung, 2003). To reflect this assumption, in the main set of simulations we only exposed our model to looming stimuli, as if STDP-driven changes were only implemented during collisions (but see sensitivity analyses below). The network was allowed to develop for 12500 time steps (at least 500 looming stimuli), and we saved its topology at five equally spaced time points during this process (Figure 7D,E), from a naive network (adjacency matrix shown in Figure 7B), to its final form (Figure 7C-E). We ran the simulation 50 times, and for each connectivity snapshot analyzed its weighted graph, as well as network responses to modeled visual stimuli: a looming stimulus, a scrambled stimulus, and a full-field flash (same as in biological experiments; Figure 7H).

The **summary of modeling results** is shown in Figure 8 (black line in each plot). In development, the network became selective for looming stimuli, both in terms of **total response** (by the end of development network responded to looming 99±9% (about 2 times) stronger than to flash; Figure 8A; “no selectivity” corresponds to the value of 0) and **average selectivity** (mean Cohen *d* of responses across all cells = 1.09±0.10; Figure 8B). The **share of cells** selective for looming stimuli also increased (Figure 8C), and plateaued at ∼98% level, as did the selectivity of the top 10% of most selective cells (Figure 8D).

**Figure 8.**
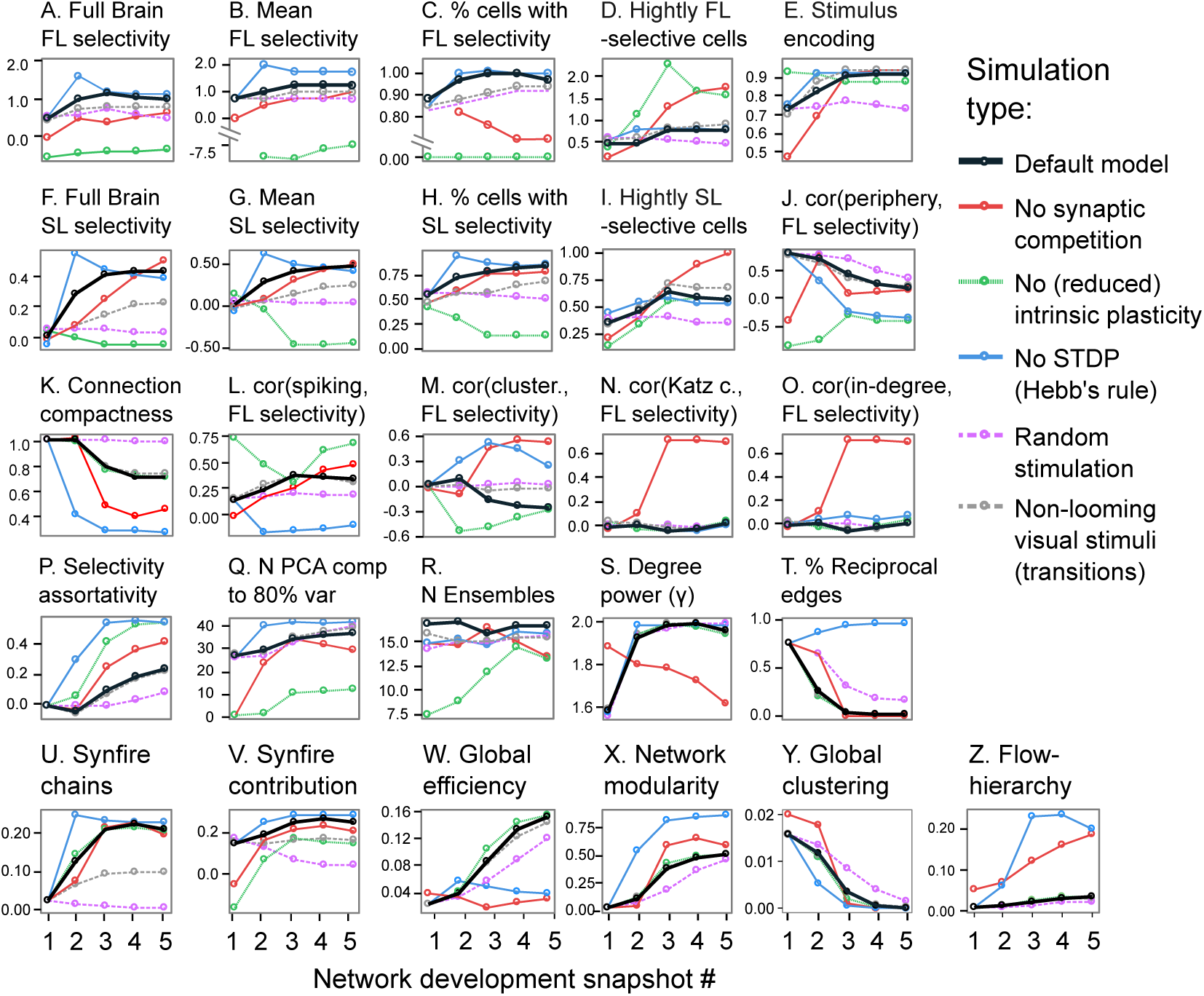
Model results. Each plot shows how one network measure changed as the model developed. Lines of different colors represent different model types: bold black is for the default model; other colors show sensitivity analyses. See the text for details. Plots B and C have a gap in Y axis to accommodate an outlier curve.

We then analyzed the quality of **stimulus encoding**, or whether stimulus identity could be reconstructed from the total activation of every cell in the network, similar to how we did it for biological experiments. For a dataset consisting of equal shares of looming and non-looming stimuli, stimulus encoding increased in development, and reached prediction accuracy of ∼95% (Figure 8E). The fact that with multivariate logistic regression we could identify looming stimuli so well suggests that a retinotopic STDP-driven network can achieve reliable looming detection, as long as it is equipped with a properly tuned output layer. We assume that in biological tadpoles, this output layer is represented by the projections from the tectum to the reticulospinal neurons in the hindbrain (Helmbrecht et al., 2018), with weights of these projections tuned in development via reinforcement learning.

Unlike for biological experiments, the model was selective for **looming over scrambled stimuli** at the full network level (Figure 8F): by the end of the training, responses to looming stimulus were 43±7% stronger than to scrambled. Mean selectivity of individual cells was 0.48±0.07 (Figure 8G); and 84% of cells were selective for looming stimuli (Figure 8H), which was also different from what we observed in real animals. Across cells within each network, looming-over-flash selectivity only weakly correlated with scrambled-over-flash selectivity (*r*= 0.17±0.10), which was unlike high correlation (r ∼0.8) observed in imaging experiments. On the contrary, looming-over-flash selectivity strongly correlated with looming-over-scramble selectivity in the model (*r*=0.91±0.02), while it was not present in imaging experiments (to r ∼0.03). This suggests that while in the biological tectum selectivity was strongly influenced by activation dynamics, in the model it was largely driven by the stimulus geometry. A subset of highly selective cells did not emerge in the model, and the difference between 90th and 50th percentiles of selectivity were rather low (∼ 0.9; Figure 8I).

The **position of selective cells** within the visual field was different in the model compared to biological experiments. While in tadpoles selective cells tended to sit in the middle of the retinotopic field, in the model they tended to be on the periphery, and selectivity for looming stimuli positively (rather than negatively as in calcium imaging experiments) correlated with the distance from the network center (Figure 8J). Similar to tadpoles, however, this correlation decreased in development (from *r* ∼ 0.75 in a naive network, to *r* ∼0.25 in a trained network). Similarly to biological networks, selective cells tended to be closer to each other (down to 71±3% of distance expected for random connections; Figure 8K), yet unlike for biological networks, this locality of connections was refined in development.

We then looked at topological and functional correlates of looming selectivity in model networks (**centrality measures**). Selective cells tended to be more spiky (*r*=0.34±0.12; Figure 8L), and usually were not a part of a cluster (Figure 8M; final correlation with local clustering coefficient *r*= −0.26±0.12). Unlike in biological networks, in the base model (red line) selective sells did not have higher Katz centrality (*r*=0.02±0.13; Figure 8N), and they did not tend to receive an unusual number of incoming connections (*r* =0.00±0.12; Figure 8O). As in biological experiments, selective cells tended to be connected to each other (weighted assortativity of 0.24±0.07; Figure 8P). In naive networks, the majority (∼85%) of strong edges (top 50% of edges by weight) tended to lead from less selective sells to more selective ones, but in developed networks this share was reduced to chance value (∼51±0.03), matching results from biological networks.

The **variability of responses** to looming stimuli over time, quantified as the number of principal components required to describe 80% of response variability, mildly increased with network development, from 28±1 to 37±1 (Figure 8Q). The number of detected network **ensembles** did not change in development, staying around 15 (Figure 8R), but the share of variance in network responses explained by the involvement of different ensembles increased from ∼35% in naive networks, to ∼50% by the end of learning. As in biological networks, cells that formed an ensemble were slightly (by ca. 20%) more likely to be connected to each other, and were about 40% closer to each other than any two random cells in the network.

The **distribution of degrees** in the model was similar to that in biological experiments: the share of weakly connected cells (weighted in-degree < 0.5) plummeted from ∼50% in naive networks to 9±2% by the end of training. On the contrary, the share of cells with degrees of 1 and 2 increased from ∼40% to 83±2%. With these changes, the power constant for the degree distribution changed from *γ* ∼ −1.5 before training to −1.96±0.03 after training for both in- and out-degrees (Figure 8S), which qualitatively matched the changes observed in biological experiments. The share of reciprocally connected cells also decreased in development (Figure 8T).

To assess whether the model developed **synfire chains** synchronized with looming stimuli, we calculated a correlation between the synaptic weights connecting different cells in the model network (coefficients of the adjacency matrix), and the frequency at which these pairs of cells were activated in a sequence (one immediately after another) during looming stimulation. We found that looming-encoding edges were overrepresented in our connectivity graphs, and their share increased in development (Figure 8U). To estimate how strongly synfire chains contributed to looming detection we looked at whether edges that led from less looming-selective to more selective cells tended to be the ones activated during looms (see Methods). We found (Figure 8V) that the contribution of looming-aligned edges to selectivity increase along the graph was non-zero, and tended to grow, but very mildly, indicating that synfire resonances were not the sole mechanism behind looming detection. Finally, we observed that most **global network measures** changed as the network matured: efficiency (Figure 8W) and modularity (Figure 8X) increased, while clustering (Figure 8Y) decreased. In all three cases, the changes were mainly due to changes in weight and degree distribution, as they persisted if calculated on networks randomized with degree-preserving rewiring. The flow hierarchy (Figure 8Z) increased mildly in development, which was entirely due to structured changes in network topology, as the effect disappeared in rewired graphs.

To conclude whether predictions of our model matched biological experiments overall, we formulated a **list of “atomic”, easily verifiable elementary statements** about different measures we analyzed (Table 1, first two columns). The model and the experiments showed similar selectivity for looming over flash, but different selectivity to looming over scrambled stimuli. The interplay between cell position and connectivity was similar, except for the spatial distribution of looming-selective cells within the retinotopic map, which was peripheral in the model, but central in tadpoles. Changes in degree distributions were well matched, yet, with a possible exception of modularity, none of other network measures matched. When correlations between different types of node centrality and cell selectivity were considered, some of them matched, but some did not.

**Table 1.** A summary of network phenomena observed in biological experiments, in comparison with simulation results for the **base** model, and several reduced models (see main text). In the table, we ✓ is used for “yes”, × for “no”, ^ for “increase”, ∨ for “decrease”, ∧∨ for “increase followed by decrease”, and = for “no change”. FL stands for “Flash-Looming” selectivity; SL - for Scrambled-Looming selectivity; “cor” abbreviates “correlation”.

| Observation | Imaging | Model |  |  |  |  |  |
| --- | --- | --- | --- | --- | --- | --- | --- |
|  |  | Base | No STDP | No Syn. Comp. | No Intrinsic | Visual | Noise |
| Full brain FL selectivity | ✓ | ✓ | ✓ | ✓ | × | ✓ | ✓ |
| Brain selectivity, change | = | ∧∨ | ∧∨ | ∧ | = | ∧ | = |
| Av. FL selectivity | 0.6 | 1.0 | 1.0 | 0.7 | −7 | 0.8 | 0.5 |
| Av. FL selectivity, change | = | ∧ | ∧∨ | ∧ | ∨∧ | ∧ | = |
| % FL select. cells | 80% | 97% | 99% | 70% | 0% | 95% | 95% |
| % FL select. cells, change | = | ∧ | ∧∨ | ∧∨ | = | ∧ | ∧ |
| Full brain SL selectivity | × | ✓ | ✓ | ✓ | × | ✓ | × |
| Av. SL selectivity | −0.1 | 0.5 | 0.4 | 0.5 | −0.4 | 0.2 | 0 |
| Av. SL selectivity, change | = | ∧ | ∧∨ | ∧ | ∨ | ∧ | = |
| % SL select. cells | 50% | 85% | 85% | 75% | 20% | 70% | 50% |
| % SL select. cells, change | = | ∧ | ∧∨ | ∧ | ∧∨ | ∧ | ∨ |
| cor(FS, FL) | ✓ | × | ✓ | ✓ | ✓ | × | × |
| cor(SL, FL) | × | ✓ | ✓ | ✓ | ✓ | ✓ | ✓ |
| Stimulus encoding, change | = | ∧ | ∧ | ∧ | ∨ | ∧ | = |
| PCA, N components | = | ∧ | ∧ | ∧∨ | ∧ | ∧ | ∧ |
| **Ensembles:** |  |  |  |  |  |  |  |
| N Ensembles, change | = | = | = | ∨ | ∧ | = | = |
| Spatial locality | ✓ | ✓ | ✓ | ✓ | ✓ | ✓ | ✓ |
| Preferential connections | ✓ | ✓ | ✓ | ✓ | ✓ | × | × |
| **Connected cells:** |  |  |  |  |  |  |  |
| Are spatially close | ✓ | ✓ | ✓ | ✓ | ✓ | ✓ | × |
| % two-way edges, change | ∨ | ∨ | ∧ | ∨ | ∨ | ∨ | ∨ |
| **Network properties:** |  |  |  |  |  |  |  |
| Degree power ($\gamma$), change | ∧ | ∧ | ∧ | ∨ | ∧ | ∧ | ∧ |
| Efficiency, change | = | ∧ | = | = | ∧ | ∧ | ∧ |
| Clustering, change | = | ∨ | ∨ | ∨ | ∨ | ∨ | ∨ |
| Modularity, change | ∧ | ∧ | ∧ | ∧ | ∧ | ∧ | ∧ |
| Hierarchy, change | = | ∧ | ∧∨ | ∧ | ∧ | ∧ | = |
| **Properties of selective cells:** |  |  |  |  |  |  |  |
| Center / Periphery location | C | P | C | P | C | P | P |
| Spatial grouping, change | ∨ | ∨ | ∨ | ∨ | ∨ | ∨ | = |
| High in-degree $k_{in}$ | ✓ | × | × | ✓ | × | × | × |
| High Katz rank | ✓ | × | × | ✓ | × | × | × |
| High activity | ✓ | ✓ | × | ✓ | ✓ | ✓ | × |
| Assortatively connected | ✓ | ✓ | ✓ | ✓ | ✓ | ✓ | ✓ |

### Sensitivity analysis

While a comparison with one faithfully constructed model is important, a better approach is to consider a family of models, and see which assumptions are critical for the replication of biological results, and which ones are not essential (Linderman and Gershman, 2017; Pauli et al., 2018). For example, we wondered whether it was important to assume that plasticity was stronger during actual collisions, or whether looming selectivity would develop if instead of looming stimuli we used more general visual stimuli. We also wondered whether structured sensory flow is necessary for the emergence of selectivity (Triplett et al., 2018). To answer these questions, we rerun our model, excluding different parts of it one by one (but not in a cumulative fashion). First we removed explicit synaptic competition, by replacing it with synapsic weight decay (Figure 8, red lines). In a different set of simulations we greatly decreased the amount of intrinsic plasticity present in the system (Figure 8, dotted green lines), and in yet another set we replaced STDP with simple symmetrical Hebbian plasticity (Figure 8, blue lines). Finally, in last two series of simulations, we let the model develop either while exposed to random visual noise (Figure 8, dashed pink lines), or a randomized mix of translating, receding, and oblique looming stimuli (Figure 8, dashed gray lines).

Disruption of individual developmental rules led to very different changes in network properties, and their trajectories. When STDP was replaced by a simple Hebbian plasticity (Table 1, column “No STDP”; blue lines in Figure 8), looming selectivity was similar to that for STDP (top lines in Figure8A,B,F,G), and the network generally developed similarly, except that reciprocal connections were not eliminated (Figure 8T); connections were more local (Figure 8K) and modular (disjointed, Figure 8X), and with very strongly interconnected ensembles (connections within ensembles were 7 times more likely than between ensembles, compared to only ∼1.2 probability ratio in base experiments).

When synaptic competition was replaced with synaptic strength decay (Table 1, column “No Syn. Comp.”), the degree distribution was very different (got flatter rather than sharper; Figure 8U, red lines); selectivity for looming stimuli was weakened (Figure 8C); the pattern of interactions between selectivity and node centrality became unlike what was observed in the base series of experiments (Figure 8M-O), and the network became strongly hierarchical (Figure 8Z). The main reason for these differences appears to be that without synaptic competition, chains of connected neurons were allowed to lead to “dead-ends” within the graph, while with competition the total output of each neuron remained constant, leading to the development of cycles.

When intrinsic plasticity was weakened (Table 1, column “No Intrinsic”; Figure 8, dotted green lines), looming detection did not emerge (bottom lines in Figure 8A-C,F-H), and networks had simpler response structure, both in terms of PCA results (Figure 8Q), and ensemble analysis (Figure 8R). This shows that agile intrinsic plasticity is critical for learning; without it the interaction between neuronal activity and network connectivity is disrupted, as the network cannot process sensory stimuli, yet is prone to spontaneous activity.

Finally, training exclusively on looming stimuli was not critical for most measurements, in terms of replicating predictions of a full model, but training on structured stimuli was. When looming stimuli (black line in Figure 8) were replaced by non-colliding visual stimuli (dashed gray line in Figure 8), most network parameters developed similarly to how they did in the base model (Table 1, column “Visual”). Stimulus encoding was similar or better; measures of looming selectivity showed similar trajectories (Figure 8A-I), except that the values of selectivity were 20-50%, and changes in many network properties were indistinguishable (Figure 8W-Z). In contrast, when patterned stimuli were replaced with uncorrelated noise (dashed pink line in Figure 8; Table 1, column “Noise”), most measures of selectivity did not improve with time (dashed pink lines are flat in Figure 8A-B and D-I), even though network topology still changed in development (Figure 8W-Z). It suggests that looming detection should not emerge in enucleated or dark-reared aquatic animals, unless the development of their tecta is guided by retinal waves with temporal statistics similar to that of behaviorally relevant visual stimuli (Huberman et al., 2008). This prediction matches experimental reports from tadpoles (Xu et al., 2011), but curiously seems to contradict observations in Zebrafish (Pietri et al., 2017).

## Discussion

In the first half of this study, we reconstruct functional connectivity in the tectum of *Xenopus* tadpoles from high-speed calcium imaging recordings, and describe novel aspects of tectal network topology. We show that tectal networks become more sparse and lightweight with development, approaching scale-free statistics (node degrees following a power distribution with *γ* > 2), and acquiring non-random network structure. We also show that looming-selective cells tend to be located in the middle of the receptive field, and that they tend to serve as “information sinks”, collecting slightly more inputs from the rest of the network, compared to non-selective cells.

In the second half of this paper, we hypothesized that a simulated developing network governed by the spike-time-dependent plasticity, synaptic competition, and stimulated by patterned visual inputs, would (1) spontaneously acquire selectivity for looming stimuli, and (2) develop a non-random network structure, (3) similar to that observed in biological experiments. The support for this hypothesis is mixed. The model did develop selectivity for looming stimuli (both in terms of average preference, and stimulus encoding), and this increase in selectivity was resilient to changes in model assumptions. Yet model result were not fully replicated in biological experiments, as in tadpoles there was no improvement in looming detection with age, as neither average selectivity, stimulus encoding, nor cell specialization increased with development. This was particularly surprising in view of a known improvement in collision avoidance in developing tadpoles (Dong et al., 2009). Moreover, while spatiotemporally tuned synfire chains clearly emerged in the model (Figure 8U), we found it hard to clearly tease out their quantitative contribution to looming selectivity.

As predicted, our model networks developed non-random structure, with a scale-free degree distribution, low clustering, and high modularity. Several results related to network structure were replicated in biological experiments (Table 1, compare columns “Imaging” and “Model / Base”): most notably, changes in network degree distribution, a decrease in bidirectional connections, and statements related to neuronal ensembles. This match between the model and the experiment suggests that at the very least, our model captured the nature of network development under the influence of synaptic competition and spike-time-dependent plasticity (STDP). Synaptic competition promoted connectivity in weakly connected neurons, while “punishing” overconnected cells, which created light-frame, openwork graph structures (Fiete et al., 2010), while STDP coordinated activity within sub-networks, increasing modularity (Stam et al., 2010; Litwin-Kumar and Doiron, 2014), similar to how it was previously described for other types of plasticity (Damicelli et al., 2018; Triplett et al., 2018). We did not observe changes in the *number* of neuronal ensembles (Avitan et al., 2017; Pietri et al., 2017), but we believe this is because our experiments were not suited for ensemble detection, as we worked with strong shared inputs that reliably activated almost every neuron in the network. This made our experiments very much unlike those that study spontaneous activity in the tectum, when different sub-networks get activated randomly, and activity propagates within modules more readily than between them (Avitan et al., 2017).

At the same time, most model predictions about how network properties were supposed to change in development did not replicate in biological experiments. There are four possible explanations for this discrepancy. First, while neurons from stage 46 and 49 tadpoles have different synaptic and intrinsic properties (Ciarleglio et al., 2015), and while retinal inputs to the tectum are known to be refined at these developmental stages (Tao and Poo, 2005; Munz et al., 2014), the patterns of internal tectal connectivity may be relatively settled by stage 46. In our model, most network measures plateaued, or even reversed late in development (Figure 8), which means that even for a qualitative comparison between the model and the experiment we have to make a critical assumption about whether developmental stage 46 corresponds to a mid-point of network maturation, or falls on the developmental plateau. The absence of improvement in stimulus encoding in older tadpoles (Figure 2G), as well as a known difference in STDP between very young (stage 42) and older (stage 48) tadpoles (Richards et al., 2010; Tsui et al., 2010), suggest that both stages included in this study may indeed fall on the “plateau”. If true, this would mean that the known improvement in collision avoidance between stage 46 and stage 49 tadpoles (Dong et al., 2009) is largely due to maturation of sensorimotor projections from the tectum to the hindbrain that we did not assess in this study.

Second, a poor fit between model predictions and biological experiments may be a consequence of a relatively low statistical power of this study. With, respectively, 14 and 16 networks reconstructed for each developmental stage, we could only hope to detect large changes in network parameters (Cohen *d* ≈ 1.0, assuming *p*_*t*_ < 0.05 threshold and 80% power). Moreover, based on available imaging studies, we can estimate that at stage 49, one half of a tadpole tectum contains about 10,000 neurons, as it measures about 40 cells across (Hiramoto and Cline, 2009), and is packed 6-10 cells deep in its thickest part (Hewapathirane et al., 2008), while tapering towards the edges (Bollmann and Engert, 2009). On the other hand, here we reconstructed connectivity within the top layer of 128±40 cells, in a field of about 12 by 12 neurons, which means that our reconstructions covered only about 1% of a full tectal network, and 0.01% of all connections. With a coverage so sparse, our parameter estimations were expected to be noisy, further lowering our test power.

Third, one can question the validity of our connectivity reconstructions, as we did not have an opportunity to compare these reconstructions to a “ground truth” connectivity (but see Xu et al. 2011). The best way to address this concern would be to run a set of control experiments, analyzing transfer entropy between pairs of cells proven to be either connected or disconnected, to estimate the power of graph reconstruction from calcium imaging recordings. Unfortunately, these experiments are beyond our technical ability, so we have to rely on indirect criteria of a successful network reconstruction. Two most important observations that support the validity of our results are the fact that the share of reciprocal connections decreased in development; and that we observed a consistent non-randomness of almost all network measurements of reconstructed networks, compared to rewired networks (Figure 5). Also, we were comforted by a good replication of tectal response shapes (Figure 2A,B, compared to Khakhalin et al. 2014), a good internal replication of edge detection between stimuli types, and an observation of retinotopy during responses to looming stimuli (Figure 3A,B).

Finally, the fourth way to explain a relatively poor fit between our model and imaging experiments is to assume that the mechanisms of looming selectivity in the tectum are after all different from that in the model. In our simulations, looming selectivity was mediated by the emergent synfire chains (Cohen-Cory, 2002; Zheng and Triesch, 2014) that were shaped by the structured sensory activation (Vislay-Meltzer et al., 2006; Clopath et al., 2010), and thus encoded activation patterns typical for looming stimuli (Pratt et al., 2008; Richards et al., 2010). When looming stimuli were presented to the model, they “resonated” with matching synfire chains, causing them to respond strongly. It may be that in the biological tectum, enhanced responses to looming stimuli are due to either delayed recurrent integration (Khakhalin et al., 2014; Jang et al., 2016), dynamic inactivation of neurons (Fotowat et al., 2011), or other non-linear effects (Baginskas and Kuras, 2009; Heap et al., 2018a). The two biggest discrepancies between the model and the experiments were the position of selective cells within the network (central in tadpoles, peripheral in the model; Figure 3C vs. Figure 8J), and the difference in signal integration in selective cells (selective cells had slightly higher in-degree and Katz rank in biological experiments, but no similar effect was observed in the base model; Figure 6A-C vs. Figure 8N,O). We however don’t find these discrepancies too problematic. The difference in spatial placement of looming-selective cells may be due to the explicit, developmentally controlled arrangement of output neurons in biological tecta that simply was not a part of our model, while the difference in Katz centrality may be explained by exaggerated synaptic competition in the model. Indeed, by introducing strong synaptic competition, we effectively forced neurons to have outputs *within* the tectal network, which led to the development of cycles, while in a real tectum selective cells may lack local outputs, projecting mostly to other brain regions. This hypothesis is supported by very high Katz centrality of looming-selective cells in simulations without synaptic competition (Figure 8N,O, red lines). We therefore believe that, with limitations of both the model and the experimental data taken into account, they were largely in agreement with each other.

At the same time, to actually use a subset of tectal cells as a functional looming detector, as we did while estimating stimulus encoding (Figure 8E), a developing brain would need access to a learning signal, to identify selective cells within the network, and wire their outputs to motor circuits. As a working hypothesis, we suggest that in aquatic vertebrates this learning signal may come from both dimming receptors in the retina (Ishikane et al., 2005; Baranauskas et al., 2012; Heap et al., 2018a), and lateral line receptors in the skin (Pratt and Aizenman, 2009; Felch et al., 2016; Truszkowski et al., 2017). These inputs can facilitate plasticity in tectal projections to the reticulospinal neurons in the hindbrain, strengthening inputs from a subset of tectal cells that were most active immediately before a collision. Moreover, during random spatial encounters, different parts of the retina would be dimmed, and different segments of the lateral line would be activated in each individual collision, allowing animals to build several overlapping subnetworks within the tectum, selective for collisions of different geometry, and projecting to different subsets of motor neurons (Frost and Sun, 2004; Barker and Baier, 2015; Helmbrecht et al., 2018). This type of learning could lead to the development of spatially nuanced escape responses that are described in both tadpoles (Khakhalin et al., 2014) and fish (Bhattacharyya et al., 2017). Based on this hypothesis, we predict that if tadpoles are raised individually in empty arenas (devoid of objects to collide with), they should have normal vision and dark-startles, but they would not be able to perform proper collision avoidance.

To sum up, we show that a combination of simple developmental rules with patterned sensory stimulation can quickly shape a random network into a structured system that is selective for looming stimuli. We show that several predictions from our model are replicated in biological experiments, although we found no improvement in looming detection between stage 46 and 49 tadpoles. We demonstrate that functional connectivity of tectal networks can be reconstructed from fast calcium imaging experiments, and that the structure of these networks can support looming detection in small aquatic animals.

## Methods

### Statistics and reporting

Unless stated otherwise, all values are reported as mean±standard deviation. For most common tests, test type is indicated by the subscript for its reported p-value: *p*_*t*_ for a two-sample t-test with two tails and unequal variances; *p*_*t*1_ for a one-sample two-tail t-test, and *p*_*r*_ for a Pearson correlation test.

When working with weight matrices, we write them as it is usually done in computational neuroscience, where *w*_*ji*_ is a weight of an edge coming from node *i* to node *j*, which is different from how adjacency matrices are presented in graph theory, where *A*_*ji*_ would typically mean an edge from node *j* to node *i* (so our **W** = **A**^T^).

### Experiments

Overall, we followed calcium imaging protocols previously described in (Xu et al., 2011; Truszkowski et al., 2017), but combined it with visual stimulation modeled after (Khakhalin et al., 2014). Experiments were performed at Brown University, in accordance with university IACUC protocols. Unless noted otherwise, chemicals were purchased from Sigma. Tadpoles were kept in Steinberg’s solution, on a 12/12 light cycle, at 18° C for 10-20 days, until they reached Nieuwkoop-Faber developmental stages of either 45-46 or 48-49. In each experiment, we anesthetized a tadpole with 0.02% tricainemethane sulfonate (MS-222) solution for 5 minutes, then paralyzed it by immersion in 20 mM solution of tubacurarine for 5 minutes, and pinned it down to a carved Sylgard block within the recording chamber filled with artificial cerebro-spinal fluid solution (ACSF: 115 mM NaCl, 4 mM KCl, 5 mM HEPES, 10 uM glycine, 10 mM glucose). The optic tectum was exposed, and ventricular membrane was removed on one side of the tectum. Tadpoles were pinned down tilted at an angle of 10-20°, to keep exposed tectal surface flat for imaging. We then surrounded the tadpole with a small circular enclosure 15 mm in diameter, made of a thicker part of a standard plastic transfer pipette, to achieve higher concentration of Ca-sensitive dye in the solution. We dissolved 50 ug of AM ester cell permeable Oregon Green 488 nm Bapta-1 (OGB1 #06807, Molecular Probes, Waltham, MA) in 30 ul of medium consisting of 4% F-127 detergent in 96% DMSO by weight; agitated this solution in a sonicator for 15 minutes, then added 30 ul of ACSF to the vial, and sonicated for another 10 minutes. The solution was then mixed with 4 ml of ACSF to the final concentration of 10 uM, transferred to the chamber, and the chamber was placed in the dark for 1 hour. After staining, the circular enclosure was removed; the preparation was gently washed 3 times, each time with 10 ml of ACSF; the chamber was filled with 10 ml of fresh ACSF, and transferred under the scope.

This staining protocol with a BAPTA-conjugated dye proved to be challenging, and had a high failure rate. As staining procedure involved a detergent, and called for high concentrations of dye, most successful preparations were those that received the highest possible exposure to the staining solution that did not yet kill the cells. A large share of preparations however either fell short of optimal staining, and had a weak fluorescence signal, and so a low signal-to-noise ratio, or got overexposed, leading to strong fluorescence, but weak responses to stimulation, as neurons grew increasingly unhealthy. This variability in signal-to-noise ratio led to differences in edge detection certainty from one experiment to another, and required a more complicated network analysis (see below).

Visual stimulation was provided with a previously described setup (Khakhalin et al., 2014), consisting of an LCD screen (Kopin Corporation, Taunton, MA, USA) illuminated by a blue LED (LXHL-LB3C, 490 nm; Lumileds Lighting, USA), with image projected to an optic multifiber (600 um, Fujikura Ltd, Tokyo, Japan). The other end of the fiber was brought to the left eye of the tadpole, and placed 400 um away from the lens, on the axis of the eye, to have the image projected to the center of the retina. The stimulation sequence consisted of three stimuli: “Looming” stimulus (in which a circle appeared in the center of the field, its radius growing linearly from nothing to a full field in 1 second), full-field “Flash”, and a spatially “Scrambled” looming stimulus. For Scrambled stimuli, we divided the field of view into a 17 by 17 grid of squares, and randomly reassigned these squares within the image. As a result, we got a stimulus that was identical to looming in terms of its total brightness at every time step, and presented fragments of a moving edge locally (within every square in a reshuffled grid), but lacked high-level spatial organization. The permutation of squares was randomized for each experiment, but was consistent between all trials within every experiment. Stimuli were delivered every 20 s, always in the same sequence (looming, flash, scrambled), typically for the total of 60 or 72 stimuli. The stimuli were generated in Matlab (Mathworks), with the help of Psychtoolbox (Kleiner et al., 2007). Excitation light for imaging was turned on one second before the onset of visual stimulation, and kept on for 5 seconds, which was previously shown not to interfere with visual responses (Xu et al., 2011).

Fluorescent responses in the tectum were imaged using a Nikon Eclipse FN1 microscope with a 60x water-immersion objective and ANDOR 860 EM-CCD camera (Andor Technologies). NIS-elements software (Nikon) was used to record the activity. We used binning with 8˗8 pixels per bin, resulting in a 130˗130 image covering the field of view of 1130 um. The data was acquired with 10 ms auto-exposure, leading to actual frame rate of 84 frames per second (11.9 ms per frame). For each preparation, we found a focal plane that produced images of as many cells as possible, which usually meant a plane focused “in-between” top-most and bottom-most cells within the field of view. To keep the signal-to-noise ratio consistent throughout the experiment despite the ongoing bleaching of the Ca sensor, we started with relatively weak illumination (with neutral density filter ND4 engaged) and no signal amplification by the camera (EM gain of 0). We then increased the EM gain level gradually after every 12 stimuli, to keep the signal level approximately constant. Once EM gain setting reached the value of 7, we increased illumination strength by disengaging one of the density filters, reduced EM gain back to 0, and repeated the process.

Videos were processed offline; circular regions of interest of equal size (21 binned pixels per region) were manually positioned over neurons with well defined Ca responses (based on the visualization of fluorescence variability in time, as generated by NIS-elements software). Average fluorescence within each region of interest was quantified for each frame, and exported to Matlab. We then processed fluorescence traces using non-negative deconvolution algorithm (Vogelstein et al., 2010), and used its output without thresholding, interpreting it as a probabilistic estimation of instantaneous spike rate for each cell. We chose not to threshold the signal, as depending on the overlap each cell body had with the focal plane, as well as the amount of dye sequestered by each cell during staining, different neurons had very different signal-to-noise ratios even within the same preparation, which complicated the matter of finding a single threshold. This decision also shaped all further steps of our analysis, as in our dataset poorly resolved cells with low spiking activity were represented not by spike traces that were mostly silent, but by traces that were noisy, and approached uniform distribution of estimator values. For each experiment, the reference cell, that was required to estimate deconvolution algorithm parameters, was selected automatically, as the cell with the fluorescent response that had the 5th highest amplitude for this preparation.

We did not attempt to match inferred spike trains to “ground-truth” electrophysiology recordings, as the validity of this calcium imaging protocol was justified previously (Xu et al., 2011; Truszkowski et al., 2017). We also did not perform background subtraction (Truszkowski et al., 2017), as most effects of background fluorescence were expected to be cancelled out during analysis (see below). The main risk of not subtracting the background is that unsubtracted traces may contain a superposition of axonal spiking and synaptic activation in the neuropil. Judging from the spatial distribution of fluorescence signals, in our experiments neuropil fibers were not stained, as the calcium sensitive dye had little access to structures below the top, exposed level of primary tectal neurons. Moreover, our signal acquisition was focused on fluorescence sources within the focal plane, meaning that any neuropil signals were both attenuated, and spatially averaged across regions of interest. Finally, neuropil activation was expected to be similar in each trial, as same stimuli were presented to the tadpole in each trial. As deconvolution operation is close to linear, and we did not perform spike thresholding, any shared neuropil signal would be deconvolved, “hidden” in inferred spike-trains, and later cancelled out during trial-reshuffling (see below). Similarly, we did not address motion artifacts, as in our preparations they were synchronous in all cells (manifested as parallel displacement of signal sources from fixed ROIs), and therefore only introduced a fixed bias to all connectivity estimations.

### Analysis

#### Basic analysis

To quantify response amplitudes, we used reconstructed responses between 250 and 2000 ms after the stimulus onset, as this window included full visual responses, but excluded artifacts caused by the excitation light. As a measure of cell selectivity, we used Cohen’s *d* effect size for the difference between responses to looming and flash, or looming and scrambled stimuli:

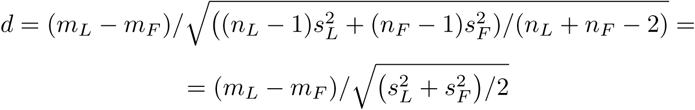

In case of equal sample sizes *n*_*L*_ = *n*_*F*_ = *n*. Here *m*_*L*_ and *m*_*F*_ are mean responses to looming and flash stimuli respectively, and *s*_*L*_, *s*_*F*_ are standard deviations for both groups. To find the **retinotopy center**, we concatenated all responses of every cell to looming stimuli into one vector, ran a principal component analysis on these vectors, then rotated two first components using promax rotation, and made sure that the 1st component *c*^1^ is the one with shorter latency, and that it is positive (by swapping and flipping the components if necessary). We then ran a non-linear optimization, looking for a pair of coordinates (*x, y*) within the field of view, that would maximize the absolute value of correlation between distances of each cell to this center and the relative prominence of the short-latency component for this cell:

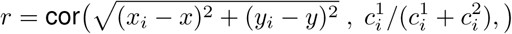

We interpreted these (*x, y*) coordinates as our best guess for the position of the “receptive field center” for each recording (the representation of the visual stimulus center within the retinotopic map). The fits were robust, with *p* < 0.05 observed in every experiment (30/30), and average achieved correlations of *r* = 0.59±0.23. To assess possible overfitting, we performed identical optimization fitting after reshuffling cell identities 5 times for each experiment, which yielded average *r* values of only 0.13±0.06, and *p*_*r*_ < 0.05 in 21% of experiments. From this we concluded that cells with early responses to looming stimuli were indeed clustered together, and that this clustering was not an artifact of our analysis, even though the *r*-values were almost certainly exaggerated due to overfitting.

For **response latency** calculations, we looked at each response *y*(*t*), and found the position of its maximum (*x*_*M*_, *y*_*M*_). We then used the least squares fit with a non-linear solver to approximate the segment between the beginning of the response and *x*_*M*_ with a piece-wise linear function:

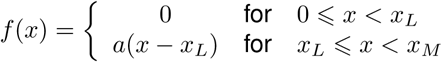

optimizing for *x*_*L*_ and *a*, where *x*_*L*_ is the response latency, and *a* is an amplitude-like parameter we did not use for subsequent analysis. This approach worked well for isolated responses with low noise, but got increasingly noisy for weak signals. Because of that, to quantify the retinotopy we used the results of factor analysis, as described above, and only used response latencies for verification.

#### Ensemble analysis

To find ensembles of cells that tended to be co-active together, we used a modified spectral clustering procedure (Ng et al., 2002) and the definitinon of spectral modularity (Newman, 2006), generalized to weighted oriented graphs. First, for each stimulus type, for each cell *i*, and separately for each experimental trial *k*, we unbiased and normalized each activity response 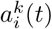, by subtracting its mean, and dividing the result over standard deviation:

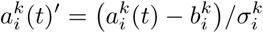

where 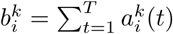and 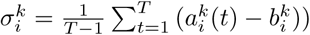.

Then, for each cell, we calculated the average response across all trials of the same type:

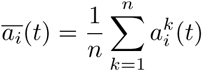

and subtracted these average responses from each trial, which resulted in a vector of a trial-by-trial deviations from the average response:

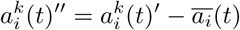

We then concatenated these vectors of deviations from the mean 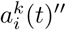 across all trials, and used them to calculate a cross-correlation matrix, to see which cells tended to be active and inactive together:.

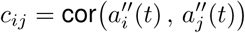

We calculated adjusted correlations *c*_*ij*_ separately for each of three types of stimuli (flash, scramble, and looming), averaged these three estimations (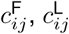, and 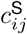) to arrive at a less noisy estimation of adjusted cross-correlation, and removed negative correlations, replacing them with zeroes:

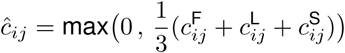

We then roughly followed the spectral clustering procedure by (Ng et al., 2002), with adjustments that seemed appropriate for ensemble detection. We first transformed our correlation matrix *ĉ*_*ij*_ into a matrix of pairwise Euclidean distances:

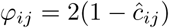

and then to affinity matrix **A**:

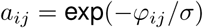

where *σ* is a free parameter that we set at 10000. We then calculated a diagonal degree matrix **D** such that *d*_*ij*_ = 0 for *i* ≠ *j*, and *d*_*ii*_ = ∑_*k*_ *a*_*i*_*k* otherwise. We used **D** to build a Laplacian matrix *L*, such that:

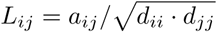

and found eigenvectors *x*_1_…*x*_*n*_ of this matrix *L*. Then we started to look for a good number of ensembles *k*, by going through all values from 1 (no ensembles) and up to the number of cells (each cell as a separate ensemble). For each *k*, we found first *k* largest eigenvectors of *L*, stacked them in columns, and renormalized each row of this matrix to give it unit length:

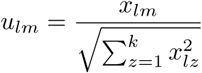

where *x*_*lm*_ is an *m*-th element of *l*-th eigenvector of *L*. We then used k-means clustering on rows of **U** as points in ℝ^*k*^, looking for *k*> clusters. Once rows of **U** (and so cells in the original data) were assigned to *k* clusters, we calculated spectral modularity of this partition on the original matrix *w*_*ij*_, using a weighted directed modification of classic formula from (Newman, 2006):

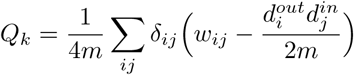

Here 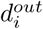 and 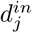 are weighted out- and in-degrees for nodes *i* and *j* respectively: 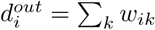, and 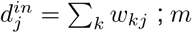 is the total strength edges involved (equivalent to the total number of edges for a non-weighted case): *m* = ∑_*ij*_ *w*_*ij*_*/*2, and *δ*_*ij*_ is a signal matrix with *δ*_*ij*_ = 1 for nodes *i* and *j* that belong to the same cluster, and *δ*_*ij*_ = 0 otherwise.

Finally, we picked the number of clusters *K* that, after spectral clustering, produced highest modularity *Q*_*K*_ across all *Q*_*k*_. We used *K* as the estimation of the number of ensembles in the network, and corresponding cluster allocation - as the allocation of cells to these ensembles.

#### Network reconstruction

For network reconstruction, we used a modified Transfer Entropy (TE) calculation, adapted from (Gourévitch and Eggermont, 2007; Stetter et al., 2012). Fast calcium imaging recordings, as used in this study, provide a middle ground between commonly used slower calcium imaging data and multielectrode recordings. In most calcium imaging recordings, the frame acquisition time (100 ms) is an order of magnitude longer than the transmission time between neurons (2 ms), which biases analysis techniques towards co-activation analyses. In our data, the high rate of video acquisition (12 ms per frame) was close to typical cell-to-cell activation transmission time in the tectum, so we restricted our analysis to interactions between the activity of each cell at a frame *t* and their activity at the next frame *t* + 1, ignoring both longer (multiframe), and same-frame interactions.

For each cell, we binned its activity trace at 3 levels, classifying every frame as either a high, medium, or low activity frame. For each cell, we used 1/3 and 2/3 quantiles of its inferred activity values as thresholds, so that all three types of frames were equally frequent, to maximize information. Then for each pair of neurons *i* and *j* we calculated the probability 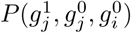, which showed the conditional probability of neuron *j* being in state 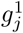(either 1, 2, or 3) at the frame *t* + 1, if this neuron was in a state 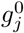 at the previous frame *t*, and input neuron *i* was in state 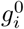 at the same frame *t*. From this set of probabilities, we calculated conditional probabilities of 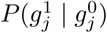, and finally calculated the total transfer entropy as

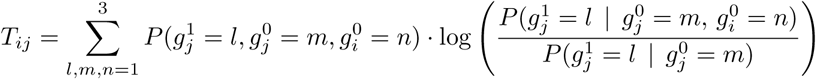

For this project, a common sensory drive (visual inputs from the retina) presented a unique problem. If the hypothesis of this paper is true, and the detection of looming stimuli in the tectum is actually mediated by sensory activation of matching synfire chains, we can expect the pattern of this sensory activation to be synchronized with causal transfer of excitation from one neuron to another. Because of that, common sensory inputs cannot be eliminated by methods that rely on the comparison of delays (Wollstadt et al., 2014). Instead, we eliminated the effects of common drive by randomly reshuffling our data within each stimulus type, and pairing activation history of each cell with activation history of other cells from non-matching trials. For each experiment, we calculated 1000 randomly reshuffled transfer entropy estimations, and then subtracted the average of these reshuffled TE estimations from raw TE estimation, arriving at adjusted TE (Gourévitch and Eggermont, 2007):

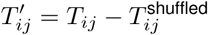

This approach is similar to the idea of analyzing subtle variations in activation from one response to another, as opposed to the analysis of activation traces themselves. As presented stimuli were same in every trial, the effect of common sensory drive over time was expected to be shared across all trials. This means that if a connection between cells *i* and *j* was suggested equally strongly by the analysis of real, and reshuffled data, these cells were sequentially driven by a common input, rather than by a true causal connection between them.

For each TE estimation, we also calculated a corresponding p-value that quantified whether actually observed TE was unusual enough (discernibly different), compared to TE estimations obtained on reshuffled data, which corresponded to a null hypothesis of no causal connections. With the computational power available to us, we could only generate 1000 surrogate reshuffled networks for every TE calculation, which made it impossible to use the false discovery rate correction, which is sometimes recommended for large-scale studies of brain connectivity (Lindner et al., 2011; Vicente et al., 2011). With ∼10^2^ neurons and 10^4^ connections the smallest possible non-zero p-value of 0.001, corresponding to finding a more extreme TE value in one out of 1000 surrogate experiments, was already larger than the Benjamini-Hochberg threshold of 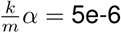. With a relatively permissive threshold of *α* = 0.01, for each of three stimuli types, our analyses suggested the existence of 5% to 69% of all possible directed edges in the connectivity graph, depending on the experiment (16% on average across experiments). As we performed edge discovery independently for responses to flashes, looms, and scrambled stimuli, we could then compare these connectivity estimations to each other. The share of edges that were suggested by responses to all three types of stimuli (on average, 0.1% of all possible edges) was 2.4 times larger, on average, than one would have expected for purely spurious discoveries with this *α* (signrank *p* = 7e-7), suggesting that the three reconstructions, originating from responses to different stimuli, could indeed be considered partial replicates for the purposes of edge discovery.

Note here that our TE adjustment procedure could not differentiate between the activity driven by shared inputs, and the activity that propagated through very reliable synaptic connections, as in both cases the same activation pattern was reproduced in every trial. It means that it was by design impossible for us to detect strongest looming-selective synfire chains in responses to looming stimuli, as the appearance of these strong connections would be indistinguishable from the effect of shared sensory input. These synfire chains could however be discovered in reponses to scrambled and flash stimuli. This means that the principle of edge replication across all three stimuli could not be taken literally, to avoid severe masking effect. Instead, we had to use a compromise approach, as described below.

Due to variations in staining quality, preparation shape, and focal plane alignment, we had to work with very different proportions of low-noise and high-noise neurons from one experiment to another, which made edge discovery rates very uneven for any fixed threshold approach. To fix this problem, we made our edge detection procedures adaptive on the experiment-by-experiment basis. First, we relaxed criteria on edge discovery, while still giving preference to edges observed in more than one subset of responses. We included in our reconstruction only edges with geometric mean of p-values below threshold: Π*p*_*k*_ < *α*^3^, where *p*_*k*_ are p-values for each of three subsets of data (responses to flash, crash, and scrambled stimuli). Then we looked for a value of *α* that would bring the average node degree (the ratio of network edges to network nodes, for directed graphs = *E/N*) to an arbitrarily set “reasonable target value”, which is a known approach to the analysis of noisy networks (Stetter et al., 2012). We picked a target *E/N* value of 1.0 (number of edges equal to the number of nodes), which lead to 128±41 edges in each experiment on average (0.9% of all possible edges); 50±21 weakly connected components, and 74±30 nodes in the largest weakly connected component. The comparison of network properties (Figures 5 and 6) did not change qualitatively in a broad range of target average degrees (*E/N* from ∼0.5 to 1.5), but observed effects became weaker and regressed to random effects outside of this range.

The TE approach does not distinguish between positive (excitatory) and negative (inhibitory) influence of one neuron on another, so our reconstructed edges could include a mix of excitatory and inhibitory connections. To estimate the share of putative inhibitory connections, we calculated pairwise correlations between activities of individual neurons, taken with a one frame delay, and compensated for the effect of shared inputs through random reshuffling, similar to how it was done for TE. We then looked at the sign of these correlations for pairs of neurons with detected connections. We found that 3±7% of detected connections seemed inactivating or inhibitory, with no difference in rate between developmental stages (*d* = 0.55, *p*_*t*_ = 0.1). According to current understanding on tectal architecture in *Xenopus*, principal tectal neurons are not expected to be inhibitory (Bell et al., 2011), and moreover, the share of negative correlations tended to be lower in experiments with better signal-to-noise ratio. We therefore assumed that most, if not all observed inactivating connections were false discoveries, and excluded all edges with negative correlation values from the analysis. For those edges that remained in the adjacency matrix, we averaged TE estimations obtained from responses to flash, scrambled, and looming stimuli, and used these averaged values as estimations of synaptic connectivity weights *w*_*ji*_.

To analyze **degree distributions**, we found the sum of weights of incoming and outgoing edges for each cell; rounded these values towards nearest whole number, and calculated frequencies *P*_*in*_(*k*) and *P*_*out*_(*k*) for each degree value *k* (Figure 4F,G). For each experiment, we then fit a regression line *y* = −*γk* + *b* to a sequence of points [*k, log*(*P* (*k*))], for in- and out-degrees separately; estimated two power constants *γ*_*in*_ and *γ*_*out*_, and averaged them to arrive at one balanced estimation (*γ*; Figure 4H).

To quantify the share of **reciprocal connections**, we multiplied the weight matrix element-wise on itself transposed, summed these values up, and normalized the result by the sum of squared weights: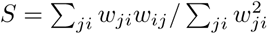. For positive weight matrices, this value is equal to 1 for symmetric matrices, 0 for matrices without reciprocal connections and smoothly changes between these two values for “intermediate” cases.

#### Network analysis

We reviewed several lists of statistical tools applicable to weighted directed graphs (Rubinov and Sporns, 2010; Costa et al., 2007; Hernández and Van Mieghem, 2011), and selected a diverse set of measures to describe different aspects of our networks, such as connectivity, unevenness of connection density, and global structure. We only included measures that do not erode with the inclusion or exclusion of individual weakly connected nodes, to make sure that metrics estimations would not change catastrophically from one experiment to another because of a slightly more generous selection of regions of interests during video quantification. Examples of measures that do not satisfy this criterion are cycle order and the “small world” property, that both are sensitive to the inclusion of weak long-ranged connections (Papo et al., 2016). We used the following list of network metrics:

#### Global network efficiency

was calculated using a function from the Brain Connectivity Toolbox (Rubinov and Sporns, 2010) on reciprocals for graph weights *R*_*ij*_ = 1*/w*_*ij*_, and was defined as:

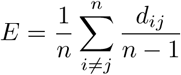

where *d*_*ij*_ is the length of the shortest path *P*_*ij*_ connecting nodes *i* and 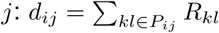

The average **clustering coefficient** (Fagiolo, 2007) was calculated using the Brain Connectivity Toolbox, with a function that supported weighted directed graphs:

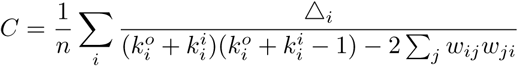

where 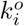 and 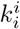 are out- and in-degrees of node *i* respectively, and Δ_*i*_ is the weighted generalization for the number of directed triangles that include node *i*:

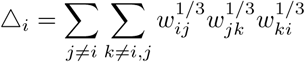

To estimate **network modularity**, we used a function from the Brain Connectivity Toolbox, which calculated spectral modularity on a weighted directed graph (Reichardt and Bornholdt, 2006; Leicht and Newman, 2008).

Our definition of **flow hierarchy** was inspired by (Mones et al., 2012; Czégel and Palla, 2015), but based on modified (weighted) Katz centrality (Katz, 1953; Fletcher and Wennekers, 2018). To calculate Katz centrality, we assumed that each node *j* collected flows of incoming signals through all edges *w*_*ji*_ leading to this node. Activation arriving through edge *j* ← *i* was proportional to the total activation *z*_*i*_ of source node *i*, the weight of this edge *w*_*ji*_, a global normalization coefficient equal to 1*/*max(*w*_*kl*_) across all edges *k* ← *l* in the network, and a damping factor of *d* = 0.9. Each node also received a small amount of constant activation (1 *− d*) = 0.1. The total activation of each node was therefore defined as:

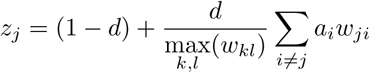

Each node then distributed this activation to other nodes. This definition is close to that of pagerank centrality (Page et al., 1999), except that the weights are not normalized to the value of total outgoing weights for each node *i*: that is, we work with raw weights of *w*_*ji*_ rather than *w*_*ji*_*/* ∑_*k*_ *w*_*ki*_. It means that a node with many outputs has a strong influence over network activation, while nodes with weak outgoing edges act as dead-ends. Similar to a standard pagerank algorithm, we solved this problem iteratively, by initializing the network with equal values of centrality, and running the equation above until convergence (typically, ∼ 100 times). Once Katz centrality values *z*_*i*_ were found, we used the difference between maximal Katz centrality observed in the network and mean centrality across all nodes as a measure of flow hierarchy: *h* = max(*z*_*i*_) − mean(*z*_*i*_) (Mones et al., 2012; Czégel and Palla, 2015).

To check whether network values described above were different from values expected on a random graph, we performed **graph randomization**, using a variant of degree-preserving reshuffling (Maslov and Sneppen, 2002), but generalized for directed weighted graphs. For a network with *N*_*E*_ edges we picked 3 · *N*_*E*_ random pairs of nodes (nodes *i, j, k*, and *l*) that had strong connections from *i* to *j*, and from *k* to *l*, but weak connections or no connections from *i* to *l*, and from *j* to *k* (we required *w*_*ji*_ > *w*_*li*_ and *w*_*lk*_ > *w*_*jk*_). We also required all four nodes to be different (*i* ≠ *j* ≠ *k* ≠ *l*). Then we cross-wired these pairs of nodes, gradually randomizing network topology:

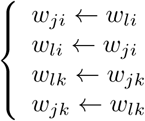

This approach to degree-preserving randomization is slightly different from the original formulation by (Maslov and Sneppen, 2002) in two ways. First, we explicitly don’t allow loops (self-edges) by requiring all four nodes be different. Second, we allow nodes *i* and *k*, as well as *l* and *j* to be connected before the rewiring, and just swap corresponding edge weights, which seems to be a necessary adjustment for directed weighted graphs. It also means that, strictly speaking, for a weighted graph, our randomization only preserves out-degrees, but not in-degrees. Because of the requirement that *w*_*ji*_ > *w*_*li*_ and *w*_*lk*_ > *w*_*jk*_, for a binary directed graph our algorithm also strictly preserves in-degrees, as it becomes identical to version by Maslov, while for nearly-binary graphs (bimodal or sparse), it tends to preserve in-degrees on average.

We also tested whether connectivity and positioning of selective cells within the graph was in any way peculiar, by calculating correlations between cell selectivity and several different **graph centrality measurements**. We used three centrality measures: weighted in-degree (a simple sum of weights of all connections to the node); Katz centrality (as described above); and node clustering coefficient (as described above, except without averaging across all nodes in the graph).

To see whether selective cells tended to form sub-networks within the graph, we calculated **weighted assortativity** of stimulus selectivity values across all nodes. The formula for the assortativity (mixing coefficient) in a weighted directed network is given in (Farine, 2014), based on the logic from (Newman, 2003) and (Leung and Chau, 2007). The original formula from (Newman, 2003) for an unweighted undirected graph defines a mixing coefficient as a Pearson correlation coefficient between properties of nodes connected by edges, taken over all edges in the graph:

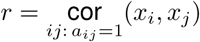

leading to the following expression:

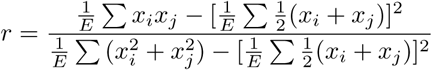

where sums are taken over all connected edges *ij*:*a*_*ij*_ = 1, and *E* is the total number of edges. For a weighted graph an equivalent measure can be introduced by replacing summation over edges to summation over all possible pairs of nodes *ij*, with weights *w*_*ij*_ introduced in each sum. The resulting expression can be rewritten in several different forms (Newman, 2003; Leung and Chau, 2007; Farine, 2014; Teller et al., 2014), but instead of explicitly coding these bulky and rather confusing calculations, we used the fact that ultimately a mixing coefficient can be described as a weighted correlation across all connected directed edges *ij* with edge values *w*_*ij*_ used as correlation weights:

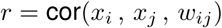

In turn, weighted correlation cor(*a, b, w*) can be introduced through weighted covariances:

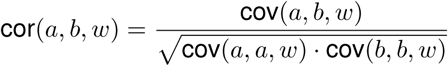

and weighted covariances are defined simply and intuitively as:

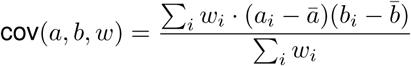

with *ā* and 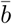 representing weighted mean values:

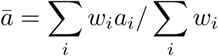

Note however that this definition may differ slightly from the one used in the Brain Connectivity Toolbox (Rubinov and Sporns, 2010).

#### Unreported analyses

For the sake of transparency, here we list all measures that were calculated, but were not included in the final manuscript for being either superfluous or confusing: four measures of weighted directed degree assortativity (in-in, in-out, out-in, and out-out); pagerank centrality; Katz centrality on reversed graphs **W**^T^; flow hierarchy for reversed graphs; node reach on both direct and reversed graphs (unweighted analog of Katz centrality without attenuation); two alternative measures of cell selectivity: McFadden’s pseudo-*R* for a logistic fit of stimulus identity to the total response of each cell, and selectivity measures calculated on peak amplitudes instead of cumulative amplitudes (the results of both calculations were qualitatively similar to those reported in the current version of the manuscript). We also made several attempts to estimate the prevalence of directed cycles in our networks, but decided that these measures require too much validation to be included in this paper. For network analysis, we also attempted to compare rewired graphs to matching random Erdos graphs, but failed to build a good generalization for a case of weighted directed graphs with an adaptive edge detection threshold.

### Developmental Model

The model consisted of *n* = 81 cells, arranged in a 9˗9 grid. The model operated in discrete time *t*, and was run for 500 epochs, 25 time steps each, or for *T* = 12500 time steps total. At each moment of time, each cell was described by three values: its instantaneous firing rate *s*_*i*_(*t*), represented as a continuous value 0 ⩽ *s*_*i*_(*t*) ⩽ 1; spiking threshold *h*_*i*_(*t*) ⩾ 0 that slowly changed over time as described below, and a constant *ŝ*_*i*_ that described the target spiking rate for each cell. Target spiking rates *ŝ*_*i*_ were randomly assigned at the beginning of each simulation, and were distributed normally around 5*/n* with a standard deviation of 1*/n*, which means that if these target spiking rates were matched, on average, at any time step, 5 out of 81 cells would be spiking.

Cells were connected to each other with “synapses” of different strengths, represented by a weight matrix **W**, with weight 0 ⩽ *w*_*ji*_ ⩽ 1. These weights were originally set at random values, uniformly distributed between 0 and 1, except for self-connections (loops, *w*_*ii*_) that were kept at 0.

At each time step we first calculated the raw activation A of all neurons: A = **W**S + B, where **W** is the connectivity matrix, S is the vector of instantaneous spiking rates *s*_*i*_, and B is the sensory input (see below). For one cell, we have:

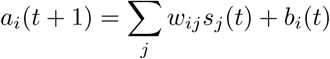

These raw activation values were then adjusted down, by a formula representing global feedback inhibition, which helped to avoid run-away excitation early in development:

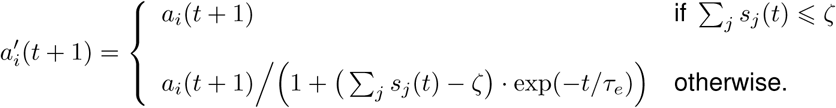

Here 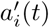 is the final, adjusted value of activation for every cell; ∑_*j*_ *s*_*j*_(*t*) is the total activity in the network at the previous time step; *ζ* is a constant that set the level of total activity at which inhibition “turns on”, and that in our case was set to *ζ* = 9 (the size of the square grid). The exponent exp(*−t/τ*_*e*_) served as an “easing” function that gradually “eased” the network from inhibition-dominated mode of operation to “free” operation, with a time constant *τ*_*e*_ = 2000. This “easing” formula was a practical compromise that greatly sped up our computational experiments, as it dampened network activity early on, when it was still close to randomly connected, and so prone to seizure-like activity, but allowed the simulation run with higher levels of activity later in development.

The activity of each neuron *s*_*i*_(*t*) was then calculated from its total activation 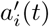:

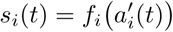

using a logistic activation function:

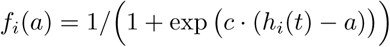

where *c* is a steepness parameter, set at *c* = 20, and *h*_*i*_(*t*) is the current spiking threshold of cell *i*. At the beginning of each simulation, spiking thresholds *h*_*i*_(0) were set to random values, uniformly distributed in a narrow band between 1*/*(*nŝ*_*i*_) and 1*/*(*nŝ*_*ii*_) + 0.1. During the simulation, the thresholds *h*_*i*_(*t*) were updated at each time step, to model the effect of **intrinsic homeostatic plasticity**. For this purpose, for each cell, we kept track of its running average spiking rate 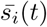, and updated both average spiking rates and spiking thresholds *h*_*i*_(*t*) by the following formulas:

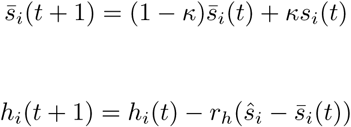

where constant *κ* = 0.05 controlled the rate of averaging, and constant *r*_*h*_ = 0.1 set the strength of homeostatic plasticity, as it controlled the rate at which spiking thresholds *h*_*i*_ were allowed to adjust, bringing the discrepancy between the target spiking rate for each cell *ŝ*_*i*_ and the running average spiking rate for this cell 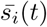 closer to zero.

Once spiking activity of each neuron at the new time step *s*_*i*_(*t*) was calculated, we moved to the **spike-time dependent plasticity** (STDP) step, adjusting synaptic weights *w*_*ji*_ that linked neurons in the network. In continuous time, STDP leads to an increase in weight *w*_*ji*_ (from *i* to *j*) if target neuron *j* spikes after a spike in source neuron *i*, with a delay that is expected for spike propagation from *i* to *j*. If the target neuron *j* spikes earlier than that (before, or together with neuron *i*), the weight *w*_*ji*_ is decreased. The amount of weight change smoothly drops off as the delay between these two spikes increases in either direction.

In discrete time, assuming that spike propagation always takes one time step, and neglecting smooth drop-off for large inter-spike intervals, STDP may be approximated by the following system of cases, with options 1 and 2 not being mutually exclusive:

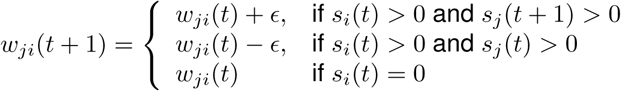

where *ϵ* is some change in synaptic weight (generally, a function of *s*_*i*_(*t*), *s*_*j*_(*t*), and *s*_*j*_(*t* + 1)).

As in our model neuronal activity *s*_*i*_(*t*) was continuous, and we wanted the synaptic change *ϵ* to be proportional to the overlap in neuronal activity, the non-exclusive system above may be rewritten as:

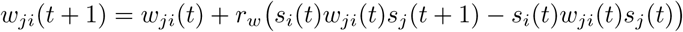

or

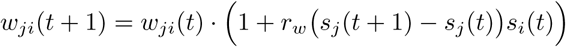

where *r*_*w*_ is a constant that controls the strength of synaptic plasticity (in this model, *r*_*w*_ = 0.25).

Finally, we modeled **synaptic competition** by introducing a negative feedback, to limit the total sum of all inputs to each neuron, and all output from each neuron. At every time step, we used the weight matrix **W** to calculate a modified matrix **W**^I^, with sums of *inputs* to each neuron normalized to a certain fixed value (set at *g* = 1.5), and a modified weight matrix **W**^o^, in which the total sum of *outputs* from each neuron was normalized to the same value:

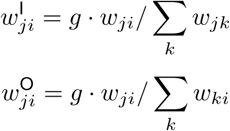

We then iteratively “moved” our actual weight matrix at each time step in the direction of the average between these two normalized matrices:

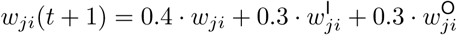

This “sliding” approach to modeling synaptic competition was less aggressive than explicit weight normalization, and allowed for more robust model convergence.

Developing networks were activated with **simulated visual stimuli** that resembled sensory activation a real animal could have experienced when navigating in a bright-lit environment with sparsely placed black spheres. For “general” visual stimulation (used in sensitivity experiments), we repeatedly created unique collision events, with randomized original positions of a black sphere relative to the eye, final distance to the eye, and direction of movement through the visual field. We would then move the projection of this virtual sphere across the virtual retina over a course of *τ* = 10 time frames. When training on looming stimuli (main series of computational experiments), we still initiated objects at randomized points within the visual field (with distances from the center of the visual field to the projection uniformly distributed between 0 and half field width) but made sure that they approached the eye on a “collision trajectory”, and eventually covered the entirety of the visual field. For looming and “general visual” stimuli, a projection of a sphere on the virtual retina was a solid circle, with its center moving linearly (*x, y*) = (*x*_0_, *y*_0_) + (*v*_*x*_, *v*_*y*_) · *t/τ*, and circle radius changing as *R*(*t*) = *R*_0_*/*(*d*_0_ − *v*_*z*_ · *t/τ*). The virtual retina consisted of 81 pixels, arranged in a 9˗9 grid (same dimensions as for the model network), with each pixel generating both “ON” and “OFF” responses without delay or bursting, based on the difference between two consecutive projections B(*t*) = XOR img(*t*), img(*t* − 1). This signal B was then inputted to matching nodes in the model network. When training on noise, we generated random noise with 9 pixels out of 81 flicking active at any given moment of time.

For testing, we compared responses to flashes, crashes, and scrambled stimuli. “Crashes” were different from looming stimuli in that the change of projection radius with time was linear 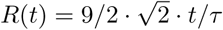, rather than realistic; this was because we used linearly expending looming stimuli in biological experiments, both in this study, and in earlier studies (Khakhalin et al., 2014). The starting position of looming stimuli varied slightly from one presentation to another (both *x* and *y* starting coordinates were normally distributed around the center of the receptive field with the standard deviation of 1 pixel), to make model responses somewhat variable, as it was necessary for our analysis. “Scrambled” stimuli were identical to “Crashes”, but with all 81 pixels randomly reassigned. “Flashes” were modeled as very fast looming stimuli that took exactly 2 frames to fill the entire field of view, and with pixels reshuffled. We used this approximation instead of a simple instantaneous flash as our model was deterministic, and we needed to introduce some variability into responses to flashes, while still keeping them as close to instantaneous as possible. As testing networks trained on different sensory stimuli, we ran into a surprising complication: during training, different stimuli provided different levels of average activation, and so not only synaptic connections between cells were differently shaped, but also intrinsic plasticity resulted in very different activation thresholds for different neurons. This variability of excitability was however an artifact of our training method, and did not approximate real biological phenomena, as for real animals, collision-like visual stimuli would happen relatively rarely, while we provided all our stimuli to the network as one intense train with no gaps. We therefore decided to let all spiking thresholds settle down before testing, to a state that was dependent only on synaptic connectivity, and not on recent stimulation history. We let the model develop for 2000 additional time steps, with only homeostatic plasticity rule on, but without STDP or synaptic scaling, while feeding neurons with Poisson random noise that activated on average 9 neurons at each time step.

The effect of this additional adjustment step was so prominent in the model, that we hypothesize that it may be indirectly relevant for the biological tectum as well. To maintain the network structure, each ensemble of synfire chains has to be regularly activated, yet the more active it is, the less excitable the neurons become, making them less likely to “win” the competition with other ensembles during stimulus classification. The dynamics of plasticity in the brain would therefore pose a meta-balancing problem (Zenke et al., 2017). If intrinsic plasticity is too fast and flexible, networks that detect unusual stimuli become overly excitable in the absence of these stimuli, producing high false-positive rates, and also habituate too fast when the stimuli are present. At the same time, due to regular activation, these networks have no trouble maintaining synaptic connections required for the stimulus detection (Litwin-Kumar and Doiron, 2014). On the other hand, if intrinsic plasticity is too slow, detection networks find it easier to maintain optimal levels of sensitivity, yet the synaptic connections that organize neurons into functional ensembles or synfire chains would erode and degrade due to disuse between stimulus presentations (Triplett et al., 2018). A potential solution to this problem is to cycle the network through distinct phases, with different contributions of synaptic and intrinsic plasticity, reminiscent of sleep-wake phases, and similar what we did in the model.

To measure the prominence of looming-resonant synfire chains in model networks we first generated a set of 100 slightly variable looming stimuli, as described above, and for each pair of cells *i* and *j* calculated the frequency at which they were activated at two consecutive time frames *L*_*ji*_. If the adjacency matrix is shaped as the imitation of sensory inputs with synfire chains (Clopath et al., 2010), we could expect *w*_*ji*_ to look like a proxy of *L*_*ji*_, while if they are not related, edges of **W** and **L** would not be correlated. To quantify that, we calculated the correlation between elements of two matrices, across all edges cor(*w*_*ji*_, *L*_*ji*_), and looked whether this correlation was higher than zero (Figure 8U). To estimate the contribution of synfire chain resonance to looming selectivity, we asked whether non-zero edges that seemed to contribute to selectivity calculations tended to be the ones activated consecutively during looms. To answer this question, we looked at a subset of edges *i* → *j* in **W** for which looming selectivity of node *j* was higher than that of node *i*, zeroed edges for which it was not the case, and calculated a correlation between elements of this modified matrix and that of the looming stimulus representation **L**. If selectivity-increasing edges within **W** are co-aligned with edges in **L**, we would expect this value of this correlation to be non-zero and positive (Figure 8V).

For **sensitivity analysis**, we either removed or greatly attenuated parts of the model, one part at a time (not cumulatively). We tried the following types of sensitivity analysis:

#### Non-looming stimuli

In this mode, instead of training the model exclusively on looming stimuli, we used a mix of randomized transitions of a black circle across the retina, as describe above. This type of stimulation was therefore still spatially patterned, but consisted mostly of translational stimuli, with some contribution of oblique looming and oblique receding stimuli.

#### Random stimulation

The network was stimulated with random noise. Each “pixel” of the image was fired with the same probability.

#### No STDP plasticity

Instead of the spike-time-dependent plasticity equation described above:

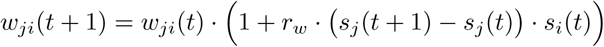

we used an equation with symmetrical Hebb plasticity, and no negative depression term:

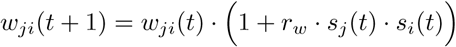

#### No synaptic competition

Instead of sliding renormalization of all inputs and outputs of each neuron, we allowed synaptic weights to slowly decay to zero: *w*_*ji*_(*t* + 1) = *w*_*ji*_(*t*) *·* (1 *− β*), where *β* = 0.001.

#### Weak homeostatic plasticity

In the formula for homeostatic plasticity, instead of change coefficient *r*_*h*_ = 0.1 we used *r*_*h*_ = 0.01.

## Acknowledgements

My greatest gratitude is to Carlos Aizenman who encouraged me to try to publish this work as a single author, even though all experiments described here were performed on his equipment, and the materials were paid for by the money from his grant (NSF IOS-1353044). I am very grateful to Lilach Avitan (The Hebrew University of Jerusalem) and Silas Busch (University of Chicago) for their in-depth reviews of the first version of this manuscript. I also thank Heng Xu (Shanghai Jiao Tong University) for his advice on imaging experiments; Joshua Vogelstein (John Hopkins) for his help with adaptive thresholding; Petko Bogdanov (SUNY Albany), Csilla Szabo (Skidmore College), Gerrit Ansmann (Bonn University), and Jim Belk (St Andrews University) for their help with network science and graph theory, and Sven Anderson (Bard College) for advice on model analysis.

The author has no conflicts of interest to disclose.

